# Sliding Window INteraction Grammar (SWING): a generalized interaction language model for peptide and protein interactions

**DOI:** 10.1101/2024.05.01.592062

**Authors:** Alisa A. Omelchenko, Jane C. Siwek, Prabal Chhibbar, Sanya Arshad, Iliyan Nazarali, Kiran Nazarali, AnnaElaine Rosengart, Javad Rahimikollu, Jeremy Tilstra, Mark J. Shlomchik, David R. Koes, Alok V. Joglekar, Jishnu Das

**Affiliations:** Center for Systems immunology, School of Medicine, University of Pittsburgh, Pittsburgh, PA, USA; Department of Immunology, School of Medicine, University of Pittsburgh, Pittsburgh, PA, USA; Department of Computational and Systems Biology, School of Medicine, University of Pittsburgh, PA, USA; The joint CMU-Pitt PhD program in computational biology, School of Medicine, University of Pittsburgh, PA, USA; Integrative systems biology PhD program, School of Medicine, University of Pittsburgh, PA, USA; Division of Rheumatology and Clinical Immunology, Department of Medicine, School of Medicine, University of Pittsburgh, PA, USA

## Abstract

The explosion of sequence data has allowed the rapid growth of protein language models (pLMs). pLMs have now been employed in many frameworks including variant-effect and peptide-specificity prediction. Traditionally, for protein-protein or peptide-protein interactions (PPIs), corresponding sequences are either co-embedded followed by post-hoc integration or the sequences are concatenated prior to embedding. Interestingly, no method utilizes a language representation of the interaction itself. We developed an interaction LM (iLM), which uses a novel language to represent interactions between protein/peptide sequences. <u>S</u>liding <u>W</u>indow <u>In</u>teraction <u>G</u>rammar (SWING) leverages differences in amino acid properties to generate an interaction vocabulary. This vocabulary is the input into a LM followed by a supervised prediction step where the LM’s representations are used as features.

SWING was first applied to predicting peptide:MHC (pMHC) interactions. SWING was not only successful at generating Class I and Class II models that have comparable prediction to state-of-the-art approaches, but the unique Mixed Class model was also successful at jointly predicting both classes. Further, the SWING model trained only on Class I alleles was predictive for Class II, a complex prediction task not attempted by any existing approach. For de novo data, using only Class I or Class II data, SWING also accurately predicted Class II pMHC interactions in murine models of SLE (MRL/lpr model) and T1D (NOD model), that were validated experimentally.

To further evaluate SWING’s generalizability, we tested its ability to predict the disruption of specific protein-protein interactions by missense mutations. Although modern methods like AlphaMissense and ESM1b can predict interfaces and variant effects/pathogenicity per mutation, they are unable to predict interaction-specific disruptions. SWING was successful at accurately predicting the impact of both Mendelian mutations and population variants on PPIs. This is the first generalizable approach that can accurately predict interaction-specific disruptions by missense mutations with only sequence information. Overall, SWING is a first-in-class generalizable zero-shot iLM that learns the language of PPIs.

## Introduction

Deep learning has revolutionized prediction across contexts^1^. In biological domains, protein language models (pLMs), have been extensively utilized for engineering synthetic protein sequences that function similarly to natural proteins^2,3^. These models have been used in a wide variety of contexts from predicting variations in human antibodies for better understanding the immune repertoire^4^ to functionally enriching the human proteome.

Deep generative models have been applied across biological contexts, including wide-ranging applications in genomics and proteomics^5,6^. Recently, AlphaFold solved the decades-old problem of protein structure prediction from sequence using evolutionary information gleaned from multiple sequence alignments^7^. ESMFold^8^ used a large language model (LLM) to leverage protein sequences and predict structures. A key conceptual difference from the AlphaFold model is that the ESMFold model circumvented the need to construct multiple sequence alignments, making downstream predictions faster. Broadly, this class of approaches, LLMs applied to proteins, are called protein language models (pLMs). They are trained to extract information from sequence, structure, or a combination thereof. However, while pLMs have been extremely successful in different contexts, they do not capture associations between proteins.

This is a critical limitation of pLMs as most proteins perform their functions via interactions with other proteins including large macromolecular complexes. Currently, pLMs have been used sub-optimally to learn the language of protein interactions. pLMs embed the sequences of the putative interactors separately and predict protein-protein interactions (PPIs) from combined embeddings^9,10^. Alternatively, they concatenate the sequences end to end and embed the concatenated sequence^11,12^. The embeddings generated typically are aggregated over the contribution of all the amino acids per position^13^, without taking into account inherent context-specific differences or the residue contact points critical to an interaction. Residue contact points are fundamental since specific residue pairs in interacting proteins are governed by evolutionary principles and are constrained^14–16^. Moreover, sequence length often presents a major obstacle in scaling pLMs to all classes of proteins.

Here we present Sliding Window Interaction Grammar (SWING), a first-in-class interaction language model (iLM) that captures the language of protein-protein and protein-peptide interactions. We applied SWING to a range of tasks, where it was comparable to or outperformed the benchmarks set by the current state-of-art methods across biological domains. We detailed the importance of our iLM in multiple nuanced interaction associated tasks such as predicting peptide-MHC (pMHC) interactions across contexts and species or the impact of missense mutations on disruption of specific protein-protein interactions. Furthermore, we have extended the utility of deep learning-based models to contexts where the full-length protein sequence is unavailable or unnecessary. Overall, SWING is a generalizable iLM that can be applied across contexts to learn the language of peptide and protein interactions.

## Results

### Overview of the SWING framework

pLMs have been widely used to predict sections of sequence that have been masked from their surrounding context in order to learn meaningful representations for multitude of tasks^8,17^. The model produces residue-level embeddings, which are aggregated to output per protein representations. However, while these models are extremely versatile at capturing individual sequence or structural information, they are sub-optimal at capturing interactions. In most current approaches, protein sequences either are concatenated and embedded or a post-hoc integration of the embeddings for the individual proteins is used for the prediction tasks^11–13^ (Fig 1A). The aggregation of per residue embedding to generate embeddings for each protein does not capture the functional importance of residues in the context of the interaction. The importance of pairwise residue interactions between two interacting proteins is well characterized^16^.

**Figure 1:**
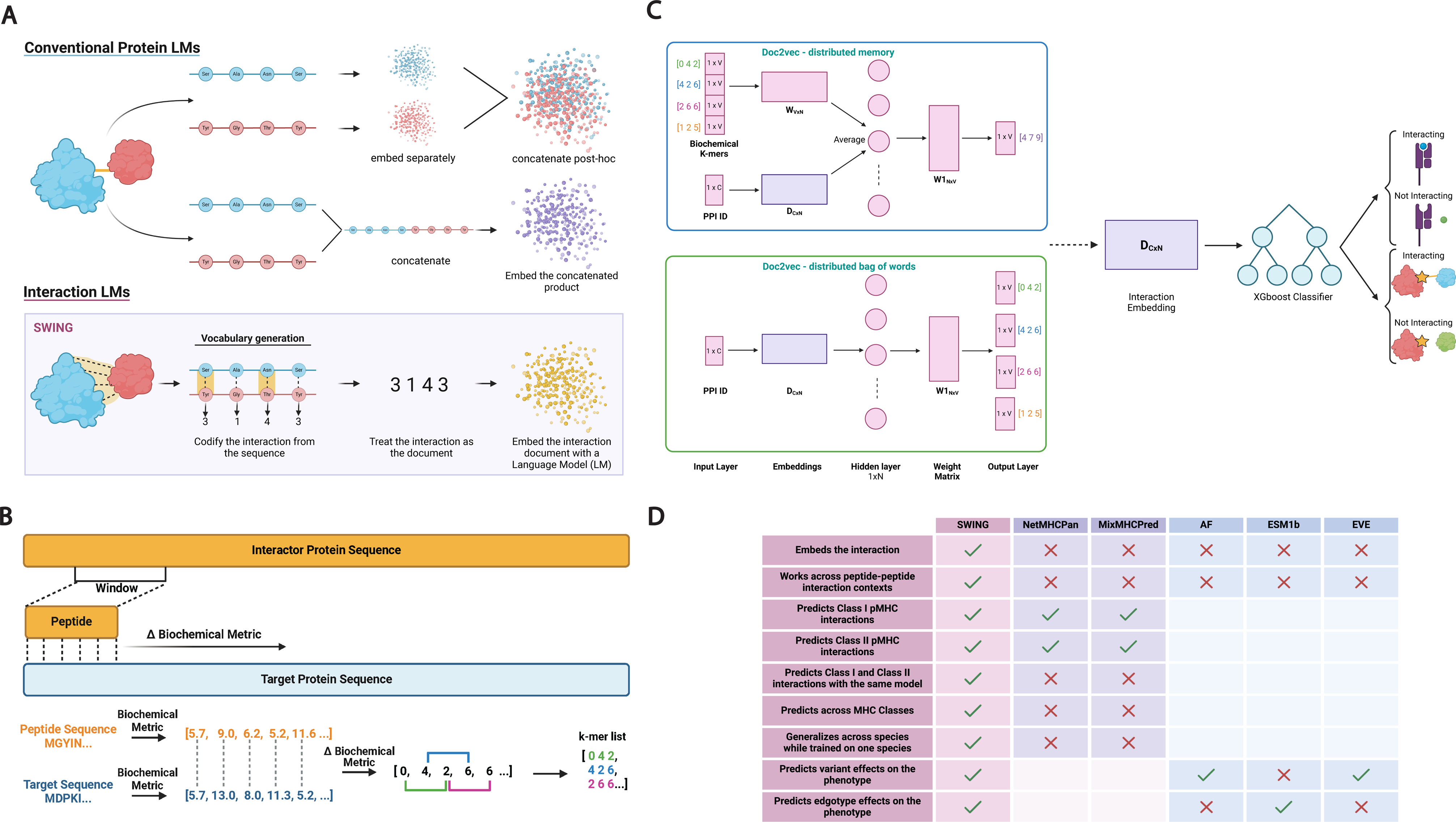
**a.** Schematic highlighting the difference between existing protein language models (pLMs) and the novelty of interaction language models (iLMs). **b.** Conceptual overview of the SWING vocabulary generation step. **c.** Abstract overview of the embedding and classification functions of SWING. Overview of Distributed memory (DM) and Distributed bag of words (DBOW) doc2vec architectures. In the doc2vec models; V is the vocabulary of the interaction language; W is the k-mers embedding matrix; C is the total number of interactions; N is the dimension of the embeddings; W1 is the output layer weight matrix **d.** Key conceptual innovations of SWING.

To address fundamental limitations of pLMs in capturing context-specific information in the context of interactions, we developed SWING, an interaction language model (iLM) that preemptively (pre-embedding) encodes pairwise residue information to generate a language-like representation (Fig 1A). The resulting sequence has encoded biochemical information and can be used for generating interaction specific embeddings without concatenation or post-hoc embedding integration. An iLM is based on capturing all the pairwise residue information in a language-like representation followed by the embedding of the interaction language. It does not rely on pLM representations for individual proteins in the interaction nor on modifying existing pLMs for interaction specific tasks.

In protein interactions, small but highly conserved sequences within each protein form the contact points between the interactors to facilitate binding^14,16^. Additionally, local information distributed within a few subsequences of a protein dictates function and structure of the whole protein^18,19^. We hypothesized that a local region in the protein of interest, determined based on the domain of application, should capture a meaningful representation within an interaction as a language. We call this subsequence the sliding window. The sliding window, a peptide of length ‘n’, is completely matched with the n positions on the target partner sequence, starting from the first position. At each position, the difference in an amino acid based biochemical metric is calculated. This difference is rounded off and the absolute value is taken to ordinally encode each amino acid pair. The use of a biochemical metric is motivated by a previous study that showed the structural importance of biochemically different amino acids^20^. Next, the sliding window is shifted by one amino acid position and the above steps are repeated until the end of the sequence (Fig 1B). We divide the resulting sequence into overlapping k-sized subsequences (k-mers), where each subsequence can be thought of as a “word” and each interaction as a “document” composed of these words. This is critical for the inherent mechanism through which embeddings are generated by the model. Importantly, this iLM architecture samples all possible biochemically informative inter-protein residue pairs to capture interaction specific information to perform many different prediction tasks.

The Doc2Vec^21^ model was used to infer the vector representation for each interaction. For the applications discussed here, we considered each interaction as a document and the k-mers as words. Further, Doc2Vec has two modes for generating the document vector - the distributed memory (DM) model and the distributed bag of words (DBOW) model. We used the different modes for different biological settings. In the DM model, each interaction and the k-mers are mapped to specific vectors. A given number of k-mers surrounding a target k-mer were considered the context of the target k-mer. Particularly, each interaction is mapped to a unique vector represented by a row in the matrix **D_CxN_**, and every k-mer is also mapped to a unique vector represented as a row in the matrix **W_VxN_**. Here, **C** is the total number of interactions and **V** is the unique set of k-mers across the interactions. The average of the embedding matrices **D** and **W** are fed into the hidden layer, which projects the averaged input vectors into a lower rank space with dimensions **N.** The output layer applies a softmax function to produce the probability distribution over the k-mers. The probability scores are used to predict the target k-mer from its context. **D** and **W** are randomly initialized with regular updates as the training progresses. In contrast to the DM model, in the DBOW model, fixed length subsequences of k-mers sampled from the interaction are defined as the context. In this architecture the interaction vector is inferred by forcing the model to predict the k-mers in the randomly sampled context from the interaction document given the interaction vector **D_CxN_**. The interaction embeddings (**D**) generated from doc2vec can then be used as features for supervised learners such as XGboost to perform various classification tasks (Fig 1C). Notably, the architecture is relatively shallow with a single hidden layer and significantly fewer parameters than LLMs. To demonstrate the applicability of these embeddings as features for wide-ranging classification tasks, we chose biologically orthogonal and representative settings. First, we focused on predicting peptide interactions to a range of Class I and Class II alleles of the major histocompatibility complex (MHC). There are significant structural differences in the formation of the pMHC complex between the two classes. MHC Class I has a deep binding groove and imposes tight constraints on the residues and peptide lengths to achieve binding. Most MHC I peptide ligands range between 8-12 residues. MHC II has shallower grooves that are comparatively open leading the N and C-terminal ends of the peptide to extend beyond the binding groove. As a result, the corresponding peptides vary in length (9-22 residues). Consequently, there is high selectivity in the binding of peptides to particular classes of MHC. Further, we applied SWING in an orthogonal context of predicting protein-protein interaction perturbation by a missense mutation. Unlike the pMHC setting, whole proteins are interacting with each other and a variant, through highly localized action, can disrupt the interaction (Fig 1C). Overall, SWING is a highly generalizable framework by design. In the subsequent sections we will systematically show that SWING can perform a range of interaction specific prediction tasks and outperforms state of art methods in different biological contexts (Fig 1D).

### SWING learns the language of peptide-MHC interactions

Classical MHC molecules are crucial in adaptive immune responses mediated by CD8 and CD4 T-cells. These T-cell immune responses are initiated by the T-cell receptor (TCR) based recognition of a peptide (antigen) bound to an MHC. MHC molecules have a highly promiscuous binding interface that allows presentation of a large number of distinct peptides, thereby displaying the cellular proteome. At a population level, MHC alleles are highly polymorphic, by virtue of which distinct alleles can present distinct repertoires of peptides. Therefore, selection of peptides to be presented by the MHC is a key selective paradigm of antigen processing and presentation^22^. Given the diversity of MHC polymorphisms (>10^4^ MHC I and >3000 MHC II alleles have been identified so far), the search space of pMHC combinations is exponentially large (MHC I: ∼10^7^ and MHC II: ∼10^20^).^23^ Overall, given the diversity of pMHC and the transient and dynamic nature of the immunopeptidome, it is impossible to experimentally identify all possible pMHC interactions^23^. Thus, a range of computational approaches have been developed to predict pMHC interactions. These include methods that try to predict pMHC binding from enriched motifs in the peptide sequence or the peptide contact points on the MHC^24^. The sequence of the MHC molecules also determines the properties of the peptides that can bind^25^ leading to approaches learning allele-specific pMHC interaction features based on experimentally determined training datasets. In addition to the sequence level diversity in the MHC, the principles of Class I and Class II binding are very different from a structural and functional perspective^26^. Existing methods do very well for alleles that they are trained on, but they often have limited efficacy for alleles for which little or no experimental data is available. They also cannot cross-predict between classes and species well.

We applied SWING to this prediction task^27^. The epitope (presented peptide) represents the essential local context within the protein sequence and is well suited as the sliding window for SWING. The functionally relevant part of the MHC sequence along with the peptide window were used to generate the biochemical language (Fig 2A) and served as the input to downstream supervised predictors. We first trained a SWING model to predict pMHC interactions for MHC I using the human immunopeptidome. Because the peptides that bind to MHC I are from endogenously synthesized proteins, this specific pMHC complex orchestrates a CD8+ T-cell response. Finally, activated CD8+ T-cells are cytotoxic and kill host cells that express antigen peptides bound to MHC I^28^. The interactions of a small number of functionally distinct human leukocyte antigens (HLAs) were chosen for training to sufficiently capture functional variability and redundancy in the dataset. We used gradient boosted trees as the supervised classifier, a standard ensemble approach that optimizes the bias-variance tradeoff through bootstrap aggregation and helps prevent overfitting (Methods).

**Figure 2:**
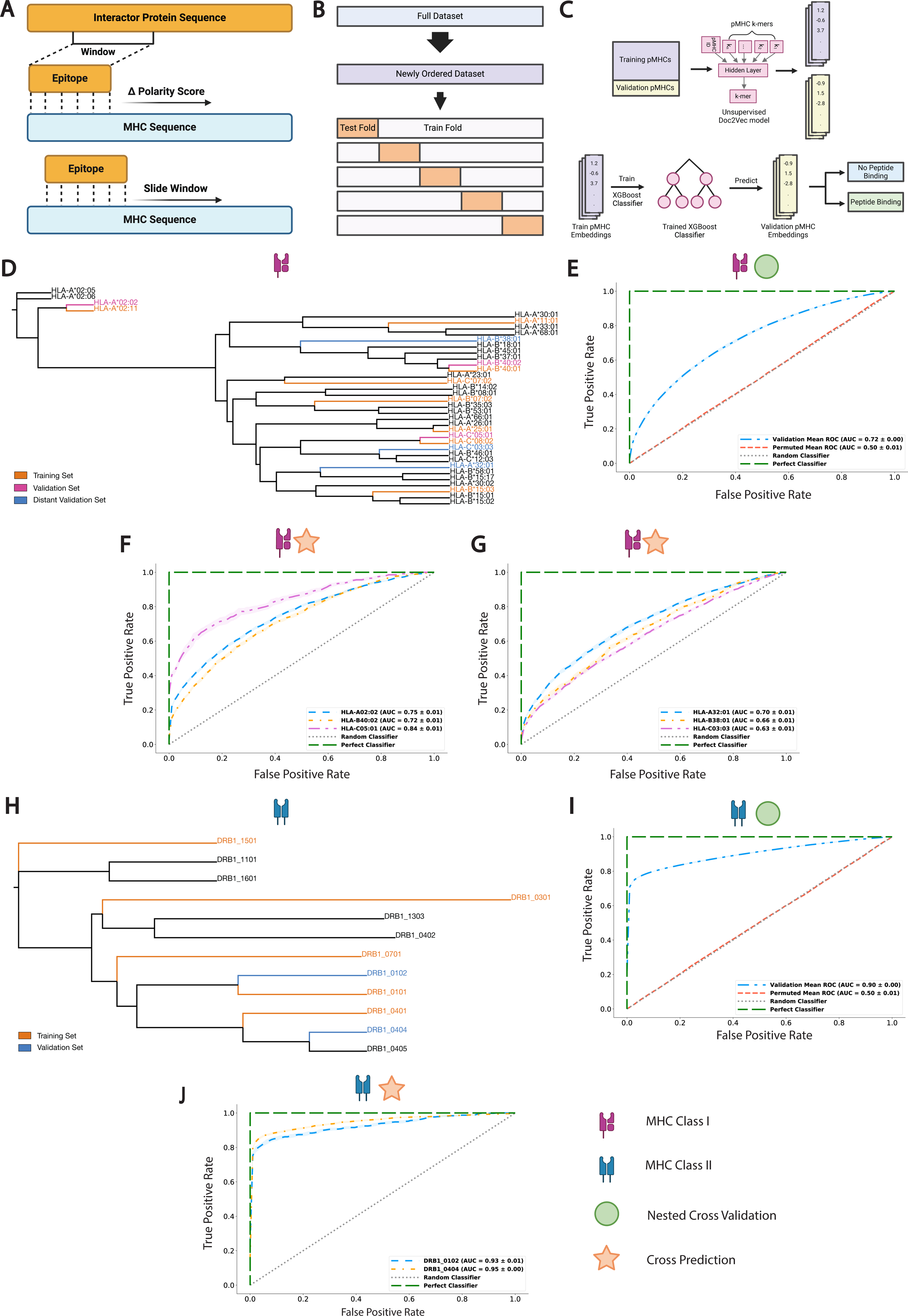
**a.** Schematic of the pMHC prediction task adaptation of the SWING framework. **b.** Representation of the standard Cross Validation (SCV) evaluation metric. **c.** Depiction of the cross-prediction evaluation metric. **d.** Class I allele functional clustering as defined by MHCcluster 2.0. Orange, alleles in the training set; magenta, alleles in the validation set; blue, distant allele validation set. **e.** SWING Class I model performance plotted across 10 replicates of 10-fold cross-validation with permutation testing defined by the area under the receiver operating characteristic curve (AUC-ROC). Blue, validation curve; red, permuted mean; green, Perfect classifier; gray, Random classifier. **f.** Class I model performance on 3 unseen functionally close alleles in the validation set as defined by the AUC-ROC. Blue, HLA-A02:02; orange, HLA-B40:02; magenta, HLA- C05:01; green, Perfect classifier; gray, Random classifier. **g.** Class I model cross-prediction performance on 3 unseen functionally distinct alleles in the distant validation set as defined by the AUC-ROC. Blue, HLA-A32:01; orange, HLA- B38:01; magenta, HLA-C03:03; green, Perfect classifier; gray, Random classifier. **h.** Class II allele functional clustering as defined by MHCcluster 2.0. Orange, alleles in the training set; blue, alleles in the validation set. **i.** Class II model performance plotted across 10 replicates of 10-fold cross-validation with permutation testing defined by the AUC-ROC. Blue, validation curve; red permuted mean; green, Perfect classifier; gray, Random classifier. **j.** Class II model cross-prediction performance on 2 unseen alleles in the validation set as defined by the AUC-ROC. Blue, DRB1_0102; orange, DRB1_0404; green, Perfect classifier; gray, Random classifier. For the AUC-ROC plots the shaded regions span +- the standard deviation.

To rigorously assess model performance, we evaluated SWING using two metrics. First, we employed cross-validation with multiple replicates of the dataset, where subsets of data are iteratively held out to later test the model (Fig 2B). This approach helps to better assess the generalizability of our model and evaluate any bias specific to the test dataset. Additionally, we performed cross-prediction (Fig 2C), where data from contexts independent of the training data are used to evaluate model performance. Here, cross validation testing datasets had alleles common to the training set, even if the two sets were mutually exclusive for pMHC interactions (Fig 2B) while for cross prediction we chose alleles that were present only in the test set (Fig 2C). Our validation tests for downstream model performance evaluation were functionally disparate from the training dataset (Fig 2D). We found that SWING was significantly predictive of peptide-MHC I binding in a k-fold cross-validation framework (Fig 2E, AUC = 0.72, P < 0.001). Additionally, we observed that prediction accuracy remains consistent for alleles absent from our training setup for all 3 alleles (Fig 2F, AUC=0.75, 0.73, 0.84 for HLA-A02:02, HLA-B40:02, and HLA-C05:01 respectively). Importantly, while the expression of specific alleles is population dependent, not every allele will be extensively sampled for its epitope profile^29^. Therefore, a key strength of SWING is its ability to predict peptides for alleles absent from its training dataset. To further cement SWING’s performance in the Class I MHC context, we chose alleles from functionally distant clusters for training and validation. Despite the difficulty of the task, SWING effectively predicted peptides for this set of unseen HLA-I alleles (Fig 2G, AUC= 0.7, 0.66, and 0.63 for HLA- A32:01, HLA-B38:01, HLA-C03:03 respectively).

Next, we extended the SWING framework and developed a model for predicting the peptides that bind MHC II molecules to mediate CD4+ T cell responses. We repeated the exercise of selecting the functionally diverse MHC II alleles as a representative dataset for MHC II (Fig 2H). SWING could confidently predict the peptide-MHC interactions for Class II alleles in a standard cross validation evaluation (Fig 2I, AUC=0.9, P < 0.001). Consistent with model performance in the Class I MHC setting, SWING also made confident predictions for 2 Class II alleles functionally different from the training set (Fig 2J, AUC= 0.93 and 0.95 for DRB1_0102 and DRB1_0404 respectively). Overall, these results demonstrate that the unique iLM architecture of SWING accurately allows us to learn the language of both Class I and Class II pMHC interactions using a small set of functionally representative alleles.

### SWING leverages biologically meaningful information to learn the language of pMHC interactions

SWING learns about different interaction contexts from specific and minimal information. We rationalize that this biological information drives SWING’s predictive power. First, we tested this notion through changing the biochemical scale for the amino acids. The biochemical metric used to codify the interaction to a language-like representation is essential to the SWING framework. While a number of biochemical effects have been shown to be major contributors in protein-protein associations^30^, polarity was chosen as the metric for applying SWING to predict pMHC interactions. We evaluated whether SWING is capturing general interaction features or those specific to polarity. We changed the metric to hydrophobicity, a distinct biochemical metric to encode the interaction language (Fig 3A). We observed that the Class I (Fig 3B, SCV AUC= 0.72, P < 0.001, cross-prediction AUC = 0.78, 0.70, and 0.81 for HLA-A02:02, HLA-B40:02, and HLA- C05:01 respectively) and Class II (Fig 3C, SCV AUC = 0.88, P < 0.001, cross prediction AUC=0.88 and 0.88 for DRB1_0102 and DRB1_0404 respectively) models accurately predicted the interactions for functionally identical and different MHC molecules when the scale was changed to capture the hydrophobic effects. Therefore, SWING captures interaction specific information that goes beyond a single type of biochemical property.

**Figure 3:**
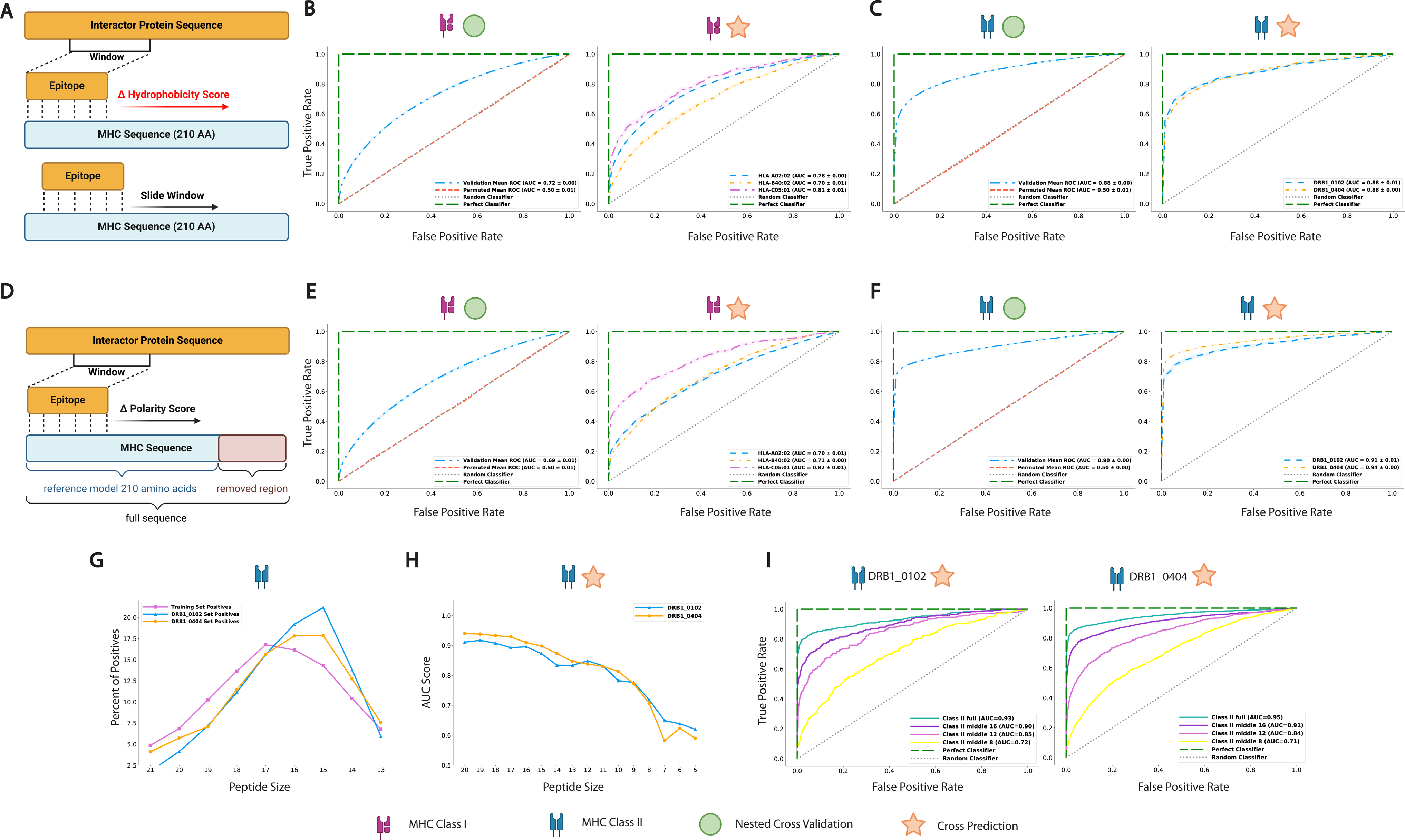
**a.** Schematic of the sequence length modification to the SWING pMHC prediction framework. **b.** Full sequence Class I model SCV performance with permutation testing defined by the AUC-ROC (left). Blue, validation curve; red, permuted mean; green, Perfect classifier; gray, Random classifier. Full sequence Class I model cross-prediction performance on 3 alleles in the validation set as defined by the AUC-ROC (right). Blue, HLA-A02:02; orange, HLA-B40:02; magenta, HLA-C05:01; green, Perfect classifier; gray, Random classifier. **c.** Full sequence Class II model SCV performance with permutation testing defined by the AUC-ROC (left). Blue, validation curve; red, permuted mean; green, Perfect classifier; gray, Random classifier. Full sequence Class II model cross-prediction performance 2 alleles in the validation set as defined by the AUC-ROC (right). Blue, DRB1_0102; orange, DRB1_0404; green, Perfect classifier; gray, Random classifier. **d.** Schematic of the hydrophobic scale modification to the SWING pMHC prediction framework. **e.** Hydrophobicity score Class I model SCV performance with permutation testing defined by the AUC-ROC (left). Blue, validation curve; red, permuted mean; green, Perfect classifier; gray, Random classifier. Hydrophobicity score Class I model cross-prediction performance on 3 alleles in the validation set as defined by the AUC-ROC (right). Blue, HLA-A02:02; orange, HLA-B40:02; magenta, HLA-C05:01; green, Perfect classifier; gray, Random classifier. **f.** Hydrophobicity score Class II model SCV performance with permutation testing defined by the AUC-ROC (left). Blue, validation curve; red, permuted mean; green, Perfect classifier; gray, Random classifier. Hydrophobicity score Class II model cross-prediction performance on 2 alleles in the validation set as defined by the AUC-ROC (right). Blue, DRB1_0102; orange, DRB1_0404; green, Perfect classifier; gray, Random classifier. **g.** Peptide length distribution of the interacting peptides in the Class II datasets defined by percentage. Magenta, training set; blue, DRB1_0102 validation set; orange, DRB1_0404 validation set. **h.** Peptide length truncation in training and test datasets affects the predictive power of the SWING Class II model as defined by the AUC for each cut-off size in 2 Class II datasets (left). Blue, DRB1_0102; orange, DRB1_0404. **i.** Visualization of the stratification of the model performance using 4 truncation cut-offs for cross predictions on DRB1_0102 (left), and DRB1_0404 (right) defined by AUC-ROC. Sea green, full length peptides; purple, 16 amino acid (AA) truncation; magenta, 12 AA truncation; yellow, 8 AA truncation. For the AUC-ROC plots the shaded regions span +- the standard deviation.

In our earlier analyses, the most relevant part of the MHC sequence was used to represent the importance of the regions participating in the peptide MHC binding. We know that the biological information pertinent to peptide binding is primarily contained in the first 206 amino acids. We assume that this part of the MHC sequence has the minimal set of information critical to the prediction task. Hence, additional domains within the sequence will not change the performance significantly. To rigorously test this assumption, we used a comparatively longer MHC sequence to test the MHC I and II models (Fig 3D). In line with expectation, we observed that the MHC length did not affect prediction quality of both Class I (Fig 3E, SCV AUC=0.69, P < 0.001, cross prediction AUC=0.70, 0.71, and 0.82 for HLA-A02:02, HLA-B40:02, and HLA-C05:01 respectively) and Class II (Fig 3F, SCV AUC=0.90, P < 0.001, cross prediction AUC = 0.91 and 0.94 for DRB1_0102 and DRB1_0404 respectively) models.

Finally, we tested the impact of biological information encoded within the peptide sequence. The core of the Class II MHC peptides is made up of 9 amino acids that determine specificity ^31,32^. Additionally, flanking peptides around the core are important for binding^33,34^. For MHC Class II, the core amino acids are found as residue motifs rather than in specific numbered positions since the peptide may slide across the MHC binding groove^35^. We analyzed the effect of removing the flanking regions and/or the core motif on SWING’s performance. We performed predictions with SWING on peptides that were truncated to up to 5 amino acids. In our training and test datasets the peptide length ranged from 13 to 21 and had a similar distribution, with most of the peptides in the range 14-18 AA (Fig 3G). The negative set in both the training and test set had an even distribution of all length peptides of ∼11% and also contained peptides of length 13-21 (Fig S1). The peptide length in both the training and test sets was reduced by one amino acid at a time while keeping the middle of the peptide. Prediction was evaluated for 2 alleles not in the Class II training set from the largest size, 21 AA, to 5 AA (Fig 3H). We observed a dramatic drop in performance once the peptide length reduced below 9 AA (Fig 3I). Consistent with biological expectation, this reflects that at least one of the amino acids, essential to the interaction, are certainly removed below 9 AA, and biological binding information is lost. While other approaches use core motifs in their learning process, SWING de-novo learns that a core motif of around 9 AAs is critical to learning the language of pMHC interactions. This confirms that SWING is not simply a black box but is learning biologically important features underlying pMHC interactions.

### SWING transfers knowledge between functionally separate interactions

The sparsity of high throughput datasets in certain biological contexts can impede discovery of relevant relationships within these complex systems. Due to the lack of training data, prediction models should be developed to learn the significantly important elements in a “data-rich” context and boost the predictions in a biologically orthogonal “data-sparse” context^36^. Few- and zero-shot learning postulate that prior knowledge acquired in one problem domain can be reused and applied to solve different but related problems^37–39^. In biology, few-shot learning and zero-shot learning have been used for context-transfer of predictive models learned in one tissue type to the distinct contexts of other tissues. These models have been used to predict phenotypic effects of gene deletions, chromatin dynamics, drug responses, and biological network dynamics in data sparse settings in precision medicine^36,40^. We rationalized that the embedded representations of biochemical motifs learned by SWING should contain interaction-specific information, which can generalize across contexts. More importantly, we wanted to test whether SWING has few-shot learning capabilities.

The ability to predict pMHC interactions across contexts is well suited to evaluate SWING’s ability to transfer interaction knowledge between multiple contexts. We chose a difficult context to evaluate SWING’s ability to make zero-shot predictions. We interrogated whether a SWING model trained on Class I pMHC interactions can learn the language of Class II pMHC interactions. There are of course key functional differences in downstream immune responses as Class I pMHC complexes signal to CD8 T cells while Class II pMHC complexes signal to CD4 T cells. In addition, there are other crucial structural differences between MHC classes. Importantly, the grooves (formed by the membrane distal domains) in MHC I molecules are closed at both ends and typically bind to peptides of length 8-12 amino acids while MHC II molecules bind peptides of 12-25 amino acids that extend beyond the ends of their open groove^41,42^ (Fig 4A). Accounting for the structural and functional differences, widely used pMHC interaction prediction tools have separate models for the two classes^43–46^.

**Figure 4:**
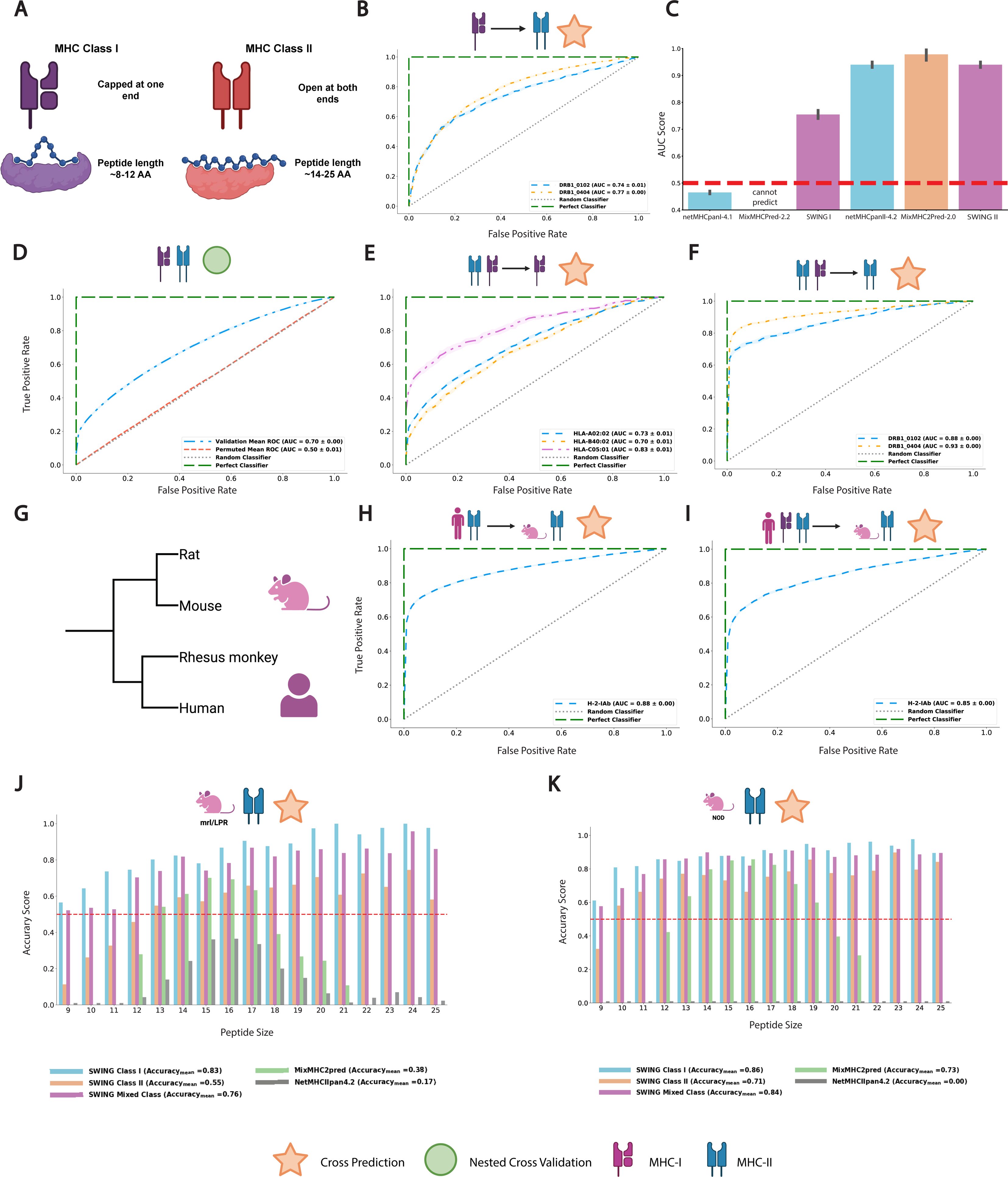
**a.** Schematic to illustrate the structural differences between the Class I and II MHC receptors. **b.** SWING Class I model performance for predicting Class II pMHC interactions defined by the AUC-ROC. Blue, DRB1_0102; orange, DRB1_0404 **c.** Comparison of AUC-ROC between SWING Class I and II models with netMHCpan 4.1, mixMHCpred 2.0, netMHCIIpan 4.2, and mixMHC2pred 2.0 models for predicting Class II pMHC interactions. Blue, netMHCpan models; orange MixMHC2Pred models; magenta SWING models. **d.** SCV prediction performance of SWING Mixed model (trained with Class I and II pMHC interactions) with permutation testing defined by the AUC-ROC. Blue, validation curve; red, permuted mean; green, perfect classifier; gray, random classifier. **e.** Prediction performance of SWING Mixed model for predicting Class I pMHC interactions represented by AUC-ROC. Blue, HLA-A02:02; orange, HLA-B40:02; magenta, HLA- C05:01; gray, Random classifier; green, Perfect Classifier. **f.** Prediction performance of SWING Mixed model for predicting Class II pMHC interactions represented by AUC-ROC. Blue, DRB1_0102; orange, DRB1_0404; gray, Random classifier; green, Perfect classifier. **g.** Schematic to illustrate the evolutionary distance between Homo sapiens and Mus musculus. **h.** Performance of the SWING human Class II model for predicting Class II pMHC interactions in mice represented by AUC-ROC. Blue, H-2-IAb; gray, Random Classifier; green, Perfect Classifier **i.** Performance of the SWING human Mixed model for predicting Class II pMHC interactions in mice represented by AUC-ROC. Blue, H-2-IAb; gray, Random Classifier; green, Perfect Classifier. **j.** Accuracy scores for H-2-IEk interacting peptides of different lengths for different SWING models, netMHCIIPan 4.2, and MixMHC2Pred 2.0. Blue, SWING Class I model; orange, SWING Class II model; magenta, SWING Mixed class model; green, MixMHC2Pred 2.0; gray, NetMHCIIpan 4.2. **k.** Accuracy scores for H-2-IAg7 interacting peptides of different lengths for different SWING models, netMHCIIPan 4.2, and MixMHC2Pred 2.0. Blue, SWING Class I model; orange, SWING Class II model; magenta, SWING Mixed model; green, MixMHC2Pred 2.0; gray, NetMHCIIpan 4.2. For the AUC-ROC plots the shaded regions span +- the standard deviation.

Despite these functional and structural differences, the SWING MHC I model for peptide binding prediction could confidently predict pMHC interactions for 2 Class II MHC molecules (Fig 4B, AUC = 0.74 and 0.77 for DRB1_0102 and DRB1_0404 respectively). Next, we compared the predictive performances of SWING in predicting Class II pMHC interactions with 2 benchmarks - netMHCpan, and MixMHCpred. NetMHCpan^24^ is a widely used tool for predicting peptide MHC interactions. Interestingly, netMHCpan4.1 and netMHCIIpan4.2 use an ensemble classifier consisting of 100 neural networks^43^ compared to the single one-hidden layer neural network architecture used in SWING for generating the interaction embeddings and gradient boosting trees for classification. MixMHC2Pred 2.0 also has a more complicated structure than SWING as it uses an ensemble of neural networks to first deconvolve the structural information and integrate sequence features where considerable changes to the architecture are made to accommodate for the nuances of each class of pMHC interaction^45^. There are also stark differences in the amount of data used for training between the models. For the Class I model, SWING uses the interactions of 8 MHC I molecules from a single species compared to netMHCpan which uses 250 MHC I molecules across multiple species. For the Class II models, the netMHCIIpan2.0 model is trained on 81 distinct MHC II alleles from multiple species and 14 unique HLA-DQ molecules. MixMHC2Pred 2.0 is trained on 88 alleles MHC II alleles which also span multiple species. However, the SWING MHC II model is trained on only 5 human Class II alleles. Although the Class II models for SWING, netMHCpan, and MixMHCpred had comparable performance for validation of Class II alleles, SWING is the only method without these alleles in its training set. Since the peptides were curated from the NetMHCIIpan4.2 training set, we show the NetMHCIIpan4.2 upper bound. Notably, the SWING MHC I model was highly predictive for Class II MHC interactions compared to netMHCpan Class I MHC model, which made random predictions for the validation sets (Fig 4C, Supplementary Table 1). Certain modifications were made to the netMHCpan Class I model in the interest of benchmarking as it is completely incompatible to run on Class II peptides. A detailed description is provided in the methods section. We were unable to use the MixMHCpred Class I model as it is incompatible with Class II data. Generally, most frameworks provide two separate models for predicting MHC I and II interacting peptides. This is acceptable due to the differences between the classes. However, Class II pMHC datasets are relatively sparse. However, the strength of our approach is that this sparsity can be alleviated by training a model on both classes of MHC, something that no existing method affords. Importantly, joint profiling of CD4 and CD8 T-cell epitopes is required to understand disease pathogenesis in multiple real world settings such as the development of vaccines against infectious agents^47^ and treatments for allergic^48^, autoimmune^49^ and neoplastic diseases^50^. The joint SWING model (trained on both Class I and II pMHC interactions) could predict correctly the pMHC interactions for data points sampled from MHC I and II molecules (Fig 4D, AUC=0.70, P < 0.001). The performance metrics were not inflated due to the presence of any one class as the performance was comparable for predicting just Class I (Fig 4E, AUCs = 0.73, 0.70 and 0.83 respectively for HLA-A02:02, HLA-B40:02, and HLA-C05:01) and Class II pMHC interactions (Fig 4F, AUCs = 0.88 and 0.93 respectively for DRB1_0102 and DRB1_0404). The fact that SWING, trained on Class I pMHC interactions is able to predict Class II MHC interacting peptides, and a joint Class I and II pMHC model can predict interactions for both classes, reflects that unique iLM architecture of SWING makes it the first method that jointly learns the shared biology of antigen binding between Class I and II.

We observed that SWING extrapolates well between the two classes while maintaining accuracy in predictions. However, the context so far was constrained to pMHC interactions in humans. However, a truly generalizable iLM should be able to predict interactions between species. To test this, we adapted the previously trained SWING models for human datasets to the problem of predicting pMHC interactions in mice. The problem also served as another test of transferring learned information in biologically distinct contexts, as mice and humans are evolutionarily distant and the MHCs for two species have widespread differences in the sequences (Fig 4G). The SWING MHC II model that was only trained on pMHC interactions of MHC II molecules in humans could predict the interactions for mouse MHC II (Fig 4H, AUC=0.88 for H-2-IAb). The joint SWING model had similar performance for the same task (Fig 4I, AUC=0.85 for H-2-IAb). We observed that the iLM SWING learns generalizable information from data in one context that are predictive in multiple different contexts. Moreover, the transfer of knowledge is specific to SWING and is not observed with widely used tools for pMHC interactions.

### SWING enables important pMHC interaction discovery in a zero-shot manner

Prior analyses demonstrate that SWING makes accurate predictions across a wide range of contexts and HLA alleles. Next, we interrogated the ability of SWING to accurately predict peptides that are eluted off of MHC in multiple disease-relevant de-novo datasets, containing peptides and alleles never seen by SWING before. This is the most stringent and perhaps the most biologically relevant validation for a new method. This has real world implications as peptide binding prediction is used to identify, rank, and select epitopes for various research directions such as immunotherapy development and vaccine design^35,51^. The peptides used by the models are restricted to 13-21 AAs with an overrepresentation of 15-17 AA peptides across training datasets. For most other methods, length bias impedes widespread discovery as key peptides may not follow structural constraints imposed by the structure of the MHC receptor and may have lengths that rarely occur in training data^52^. To that end, we focused on the mouse H-2-IEk allele, which is one of the two Class II MHC alleles in a murine model of Lupus nephritis, MRL/lpr mice. As MRL/lpr mice recapitulate several features of SLE, including infiltration of immune cells into kidneys, we interrogated if SWING would be able to accurately predict the immunopeptidome of these kidneys. The corresponding immunopeptides of female MRL/lpr mice were experimentally characterized and binders of H-2-IEk were subsequently analyzed using SWING as well as two benchmark approaches - MixMHC2Pred2.0 and NetMHCIIPan 4.2. All three SWING models (built using only Class I or Class II data and the Mixed Class model) performed significantly better than the benchmarks despite being the only method without this allele in the training data (Fig 4J, Supplementary Table 2). Of the SWING models, the Class I and mixed models performed even better than the Class II model itself. This can be attributed to the limited functional cluster distance and availability of single Class II allele datasets. More importantly, it demonstrates that SWING learns general principles to make accurate zero-shot predictions in otherwise data-poor settings. Notably, the accuracy separated by peptide length was consistent across the range only for SWING while NetMHCPan had the lowest accuracy across the different peptide lengths. MixMHC2Pred completely missed all peptides outside its working range of 12-21 AA and SWING models alone had predictive power for peptides larger than 18 AA. We observed a clear bias in performance for MixMHC2Pred and netMHCPan, where the more abundant peptides of certain lengths had higher performance. Unlike these methods, SWING predicted more interacting peptides at a uniformly high accuracy across most peptide lengths.

Often, rare variants and alleles of interest fall outside the scope of curation techniques for different functional datasets. Researchers use the mentioned in-silico predictive methods (NetMHCPan, MixMHCPred) to identify which peptides are candidates for their MHC molecule. We identified an allele, H-2-IAg7, that is not represented in the NetMHCIIPan dataset to assess how the methods compare for a custom MHC sequence input. H-2-IAg7 is a Class II mouse allele of Non-Obese Diabetic (NOD) mice that shares physiological similarity with HLA-DQ8, a high-risk allele for Type 1 Diabetes. We have previously used an immunopeptidome dataset from I-Ag7^53^ to identify autoantigens in Type 1 Diabetes^54^. I-Ag7 was also not present in the training dataset of SWING. Again, SWING models trained on Class I, II, and a mixture of the two classes reported the highest number of peptides as interacting (Fig 4K, Supplementary Table 2). MixMHC2Pred2.0 accuracy was comparably worse than the Class I and Mixed Class model even though the accuracy was only calculated for the 12-21 AA range. This is notable as the predictions by SWING were made through transfer learning whereas mixMHC2pred is specifically a Class II model with H-2-IAg7 in its training set. Surprisingly, NetMHCIIpan4.2 failed to identify a single epitope as a binder even though it incorporates various mouse alleles in its training. NetMHCIIpan4.2 returned the default lowest rank for all peptides, showing that it is unable to rank peptides from this receptor. SWING’s unique framework is unencumbered by the necessity of unique datasets and succeeds at identifying binders regardless of MHC receptor sequence, species, or peptide length. These analyses demonstrate the capability of SWING to identify de-novo biologically meaningful interactions from datasets generated for particular diseases because of its unique training setup and knowledge transferability.

### SWING predicts protein interaction perturbation by missense variants

SWING is highly adaptable to different contexts. To demonstrate this, we applied SWING to a completely different task from predicting pMHC interactions. Although millions of coding variants have been discovered in humans, most of them are variants of unknown significance. Of these, missense variants are coding variants that cause a point substitution in the amino acid sequence. Each substitution has the potential to affect protein function. A large number of computational approaches have focused on distinguishing between pathogenic and non-pathogenic missense variants by incorporating prior knowledge pertaining to mutation-associated biophysical, biochemical and evolutionary constraints. The primary focus has been on rare variants which disrupt protein structure or affect protein abundance. Hence, earlier pathogenicity prediction efforts have relied on conservation statistics, sequence and structural features^55–57^. Recently, deep learning methods have been very successful in predicting the effect of variants with high precision. EVE captures evolutionary information in biological sequences as embeddings for variant effect prediction^58^. AlphaMissense and ESM1b are large language models that can model proteins as a language by learning amino acid distributions conditioned on the sequence context^59,60^. However, most missense variants, disease-causing or otherwise, do not have an impact on the overall structure of the protein, but rather disrupt specific protein-protein interactions^61–63^. These variants are known as edgetic variants, and the corresponding interaction-specific molecular phenotypes are “edgotypes”. Edgotypes present a fundamental link between genotype and phenotype^64^, but the development of tools for their discovery remains significantly limited. Structural priors such as the localization of a variant at the binding interface of proteins has been effective in prioritizing edgetic variants. Language model-based approaches can also make distinctions between interface and not interface variants (Figs 5A and 5B). However, even state-of-the-art variant effect predictors including AlphaMissense, ESM1b, and EVE were not at all predictive of interaction-specific disruptions (Fig 5C). This continues to represent a key unmet need.

**Figure 5:**
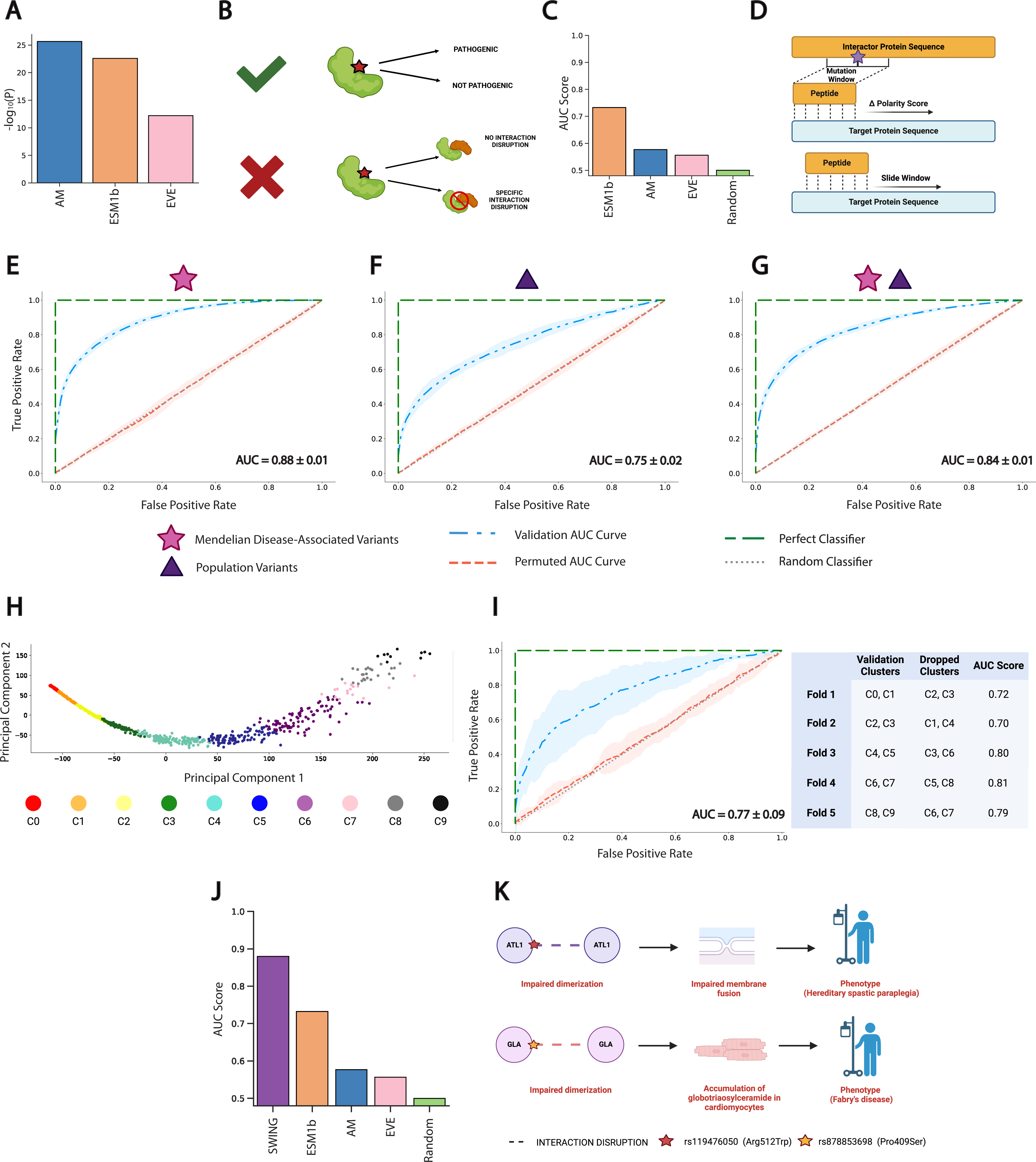
**a.** Comparison of interface variants and non-interface variants scores from the state-of-art variant effect prediction (VEP) tools represented as the negative log of the p value. A Mann-Whitney U test was performed for testing statistical significance. Blue, AlphaMissense; Orange, ESM1b; Pink, EVE **b.** Schematic illustrating the focus of current VEP tools on predicting organismal level pathogenic effect of variants over interaction specific effect. **c.** Comparison of predictive performance of various VEP tools in predicting interaction specific effects of the variants. Orange, ESM1b; Blue, AlphaMissense; Pink, EVE; Green, Random. **d.** Schematic explaining the interaction language generation procedure for applying the SWING model to predict the effect of mutations on protein-protein interactions. **e.** Prediction performance in a standard cross validation setting of SWING trained for predicting the interaction perturbation effect by mendelian missense mutations. Blue, Mendelian disease associated interaction perturbation effects; Red, Permuted classifier; Grey, Random Classifier; Green, Perfect classifier. **f.** Prediction performance in a standard cross validation setting of SWING trained for predicting the interaction perturbation effect by population variants. Blue, Population variants; Red, Permuted classifier; Grey, Random Classifier; Green, Perfect classifier. **g.** Prediction performance in a standard cross validation setting of SWING trained for predicting the interaction perturbation effect by population variants and mendelian missense mutations (Mixed). Blue, Mixed variant set; Red, Permuted classifier; Grey, Random Classifier; Green, Perfect classifier. **h.** Clusters of sequences of variant targeted proteins for the mixed dataset. **i.** Prediction performance of SWING on predicting the left-out sequence cluster represented through an AUC-ROC. Blue, Validation clusters; Red, Permuted classifier; Grey, Random Classifier; Green, Perfect classifier. The table provides the AUC for each test cluster and the corresponding proximal clusters held out from training. **j.** Comparison of predictive performance of various VEP tools in predicting interaction specific effects of the variants and SWING. Purple, SWING; Orange, ESM1b; Blue, AlphaMissense; Pink, EVE; Green, Random. **k.** Schematic representing predicted candidate interaction disrupting variants and the possible downstream effects of these in particular disorders. For the AUC-ROC plots the shaded regions span +- the standard deviation.

SWING was adapted to predict interaction specific variant effects from sequence alone. We hypothesized that the local context around the missense substitution will be the most informative of the interaction disruption phenotypes^65^. Therefore, the sliding window was selected as a subsequence of the ‘n’ amino acids in both directions of the position where the amino acid has been substituted. The parameter n was tuned based on the model performance during cross validation. The interaction partner full length sequence was the base for the target protein subsequence window to slide on to generate the biochemical enriched language for this context (Fig 5D).

First, SWING was applied to predict the effect of mutations on PPIs across various human Mendelian disorders. The data was generated from efforts to systematically characterize the human disease variome in the context of the disruptions of specific protein-protein interactions^63^. Using this dataset, in a stringent cross-validation framework, we observed that SWING could predict the effect of Mendelian disease associated disruptions with high accuracy (Fig 5E, SCV AUC=0.88, P < 0.001). Our prior work^61^ has also shown that there are interaction-disruptive single nucleotide variants (SNVs) across the allele frequency spectrum, with disruption rate being inversely proportional to allele frequency. SWING was also predictive of interaction disruption driven by population variants across the allele frequency spectrum (Fig 5F, SCV AUC=0.75, P<0.001). Together, these results show that SWING can accurately predict, relative to experimental benchmarks, the impact of missense variants on corresponding protein-protein interactions.

Next, we wanted to evaluate the feasibility of having a universal model for predicting the interaction perturbation effect of a variant, irrespective of its source (Mendelian or SNV). SWING was effective in predicting interaction disruptions caused by variants regardless of the context, in a rigorous cross-validation framework (Fig 5G, SCV AUC=0.84, P<0.001). Consistent with our observations in predicting pMHC interactions, the change of biochemical scale from polarity to hydrophobicity did not make a difference in SWING performance (Supplementary Figure 2, SCV AUC=0.85, P<0.001; SCV AUC=0.73, P<0.001; SCV AUC=0.80, P<0.001 respectively).

Furthermore, we tested whether embeddings are the more important factor in ensuring robust predictions across contexts, or whether the specific classification model used on the embeddings) were a driver of model performance. We observed that the use of logistic regression, a simpler model than XGBoost, gave us a similar performance outcome (Supplementary Figure 3, SCV AUC=0.77, P<0.001; SCV AUC=0.63, P<0.001; SCV AUC=0.71, P<0.001 respectively) and this was also the case when a more complex multi-layer perceptron was applied jointly with the embeddings (Supplementary Figure 4, SCV AUC=0.87, P<0.001; SCV AUC=0.71, P<0.001; SCV AUC=0.83, P<0.001 respectively) across variant contexts. These results demonstrate that the embeddings represent the unique iLM architecture of SWING, rather than the downstream classifier drive model performance.

As machine learning models have been shown to be able to perform variant effect prediction from sequence similarity alone^59,60^, we wanted to evaluate the impact of sequence similarity in model performance. Therefore, we divided the dataset into sequence-based clusters (Fig 5H). Each cluster was considered a fold and the test set had distinct proteins from the training set. Furthermore, we dropped the clusters adjacent to the test cluster from our training set to evaluate how the model performs when proteins similar in sequence to the test set are completely removed from the training set. We observed robust prediction performance across the different sequence clusters (Fig 5I, SCV AUC=0.77, P<0.005). The performance is similar to the models discussed above, demonstrating that the performance of SWING is primarily due to it learning higher-order sequence information beyond similarity. Overall, SWING significantly outperformed state-of-the-art VEP models including EVE, AlphaMissense, and ESM1b (Fig 5J). This is despite the fact these VEP methods have a number of parameters orders of magnitude higher than the shallow doc2vec architecture used in the SWING framework.

Cheng et al^59^. ran AlphaMissense on a set of ClinVar variants to benchmark the method against known and studied variant pathogenicity. Interestingly, several of these ClinVar variants were predicted to be benign by AlphaMissense but SWING predicted them as disruptive of corresponding protein interactions (Fig 5K, Supplementary Table 3). We focused on a couple of these variants to reconcile the potential discrepancy between AlphaMissense and SWING. The first variant was at the interface of the ATL1-ATL1 complex. This particular rare variant (rs119476050) has been reported in patients with autosomal dominant hereditary spastic paraplegia and is hypothesized to effect the protein function of ATL1^66,67^. ATL1 is important for the maintenance of the endoplasmic reticulum through membrane fusion. Importantly, dimerization of ATL1 is critical to drive membrane fusion through an enzymatic action through nucleotide hydrolysis and conformational reorganization. The monomeric ATL1 does not have enzymatic activity^68^. Furthermore, this particular variant had been cited as “likely pathogenic” in ClinVar based on the clinical evidence reported for the mutation. This represents a clear instance of a false negative of AlphaMissense that is correctly classified as disruptive of the corresponding protein-protein interaction by SWING.

Similarly, we found a mutation (rs878853698) at the interface of the GLA-GLA complex. This gene has been reported to be associated with Fabry disease. The lysosomal enzyme alpha-galactosidase A is encoded by GLA. The deficiency of this enzyme leads to the accumulation of globotriaosylceramide within cardiac myocytes leading to hypertrophy, fibrosis, apoptosis and necrosis^69^. The reported variant was found to have a damaging effect to the enzymatic activity of alpha-galactosidase A in a patient with fabry’s disease^70^. Once transported to the lysosome, GLA undergoes dimerization to become functional. It has been shown that amino acid changes associated with GLA in Fabry’s disease can lead to a fraction of the wild type levels of functional homodimers with lower or no function with an increase in the presence of heterodimers^71^. Additionally, based on the clinical evidence, this variant was also marked as “likely pathogenic” by ClinVar and missed by AlphaMissense.

## Discussion

Protein language models (pLMs) have become critical across a range of application contexts spanning protein structure prediction and engineering of novel peptides for applications in synthetic biology. However, despite their versatility, pLMs do not capture the nuances of protein/peptide interactions. We present SWING, a first-in-class interaction language model (iLM) for converting interacting proteins/peptides into a unified sequential language enriched with biochemical motifs. The language was embedded through a doc2vec architecture. Doc2vec utilizes many fold lower parameters than the architectures currently in use as protein language models. The framework is inspired from an important perspective of treating macromolecular interactions as individual entities and not as an aggregation of their constituting parts. This is seen in functional differences between dimers and monomers along with dedicated efforts to consider protein interactions as drug targets over individual proteins. Biochemical motifs are captured in the doc2vec embeddings, which were shown to be useful in varied tasks across biological domains.

The first context we applied SWING to was peptide-MHC interaction prediction. Complex formation is critical for the presentation of the antigen and upstream of robust T-cell response. This is a cornerstone of the adaptive immune system involved in surveying the host proteome for undesirable intruders. SWING was highly predictive of pMHC interactions across alleles, contexts (Class I to Class II) and species (human and mouse). While state-of-the-art pMHC binding prediction methods train on a large number of alleles and have Class I or Class II-specific models, SWING trained only on a small number of alleles and learnt the language of pMHC interactions across Class I and Class II, as well as human and mice. Most importantly, it was able to make accurate zero-shot predictions for disease-relevant alleles in SLE and T1D that the model had never seen before. The unique iLM architecture of SWING enables these accurate predictions. While other approaches make allele-specific predictions using allele-specific models, SWING is a first-in-class approach that learns the shared basis of Class I and Class II pMHC interactions across human and mice from a small number of representative human alleles. The ability of SWING to make zero-shot predictions and transfer knowledge across organisms reflects its robustness and versatility.

Next, we adapted SWING to perform a nuanced variant effect prediction task. The effect of mutations is a multiway setting in proteins, where most mutations take the route of altering the interaction profile rather than disrupting the structure of the target protein. However, the state-of-the-art deep learning methods (AlphaMissense, ESM1b, and EVE) only predict pathogenicity at the level of organismal fitness and not at the molecular phenotype resolution. SWING was successful at accurately predicting the impact of mendelian mutations and population variants on protein-protein interactions. Further, in a de novo setting, SWING picked up disease associated pathogenic variants at the interface of functionally important dimers, which were predicted to be benign by AlphaMissense. This is the first generalizable approach that can accurately predict interaction specific disruptions by missense mutations with only sequence information.

Diverse molecular networks drive basic biological processes and their perturbations are involved in disease pathogenesis. Massive multi-omic profiling is needed at different resolutions to understand the breadths of these complex systems. Therefore, most networks remain poorly profiled. The individual components of the SWING framework are designed to depend on easily available features such as sequences and biologically informative metrics. We have focused on predicting protein/peptide interactions, but SWING can be generalized to various macromolecular interactions. In the protein context, further nuance could be captured in PPIs, such as the effect of post-translational modifications on the interaction profile of a protein. Beyond protein contexts, SWING could be applied to gene regulatory networks, where different biological entities interact to control gene expression in a cell-type specific manner. SWING can potentially predict the RNA-RNA, RNA-DNA, protein-RNA and protein-DNA interactions in such networks. SWING prediction tasks can also be adjusted based on the context. Interaction specific tasks at different granularities such as prediction of binding affinity, antigen immunogenicity, patient outcome, etc, can also be modeled using SWING.

pLMs are foundation models which have been trained on over millions of proteins across thousands of families. Immense data sufficiently trains highly parameterized transformers, which can adapt to multiple different proteomic prediction tasks through self-supervised learning without training the model from scratch. Although SWING is generalizable across different interaction specific tasks within a biological domain, it is not a biological interactions foundation model. SWING needs to be customized and trained in a biological context dependent manner. A major reason is incompleteness inherent in biological networks as compared to proteins and the specificity of the interaction partners is conditioned strongly on the context.

Overall SWING is a highly versatile framework that can be adopted across domains for different prediction tasks. It is adept at learning the language of interactions, which transfers well as features across biological contexts within a domain and proves to be an effective tool even in data-sparse contexts.

## Methods

### SWING framework

Language generation for an interaction has three crucial components: a sliding window, the target protein sequence, and a biochemical metric. The sliding window is a subsequence from the protein of interest. The length of the sliding window can span from 3 amino acids to 25 amino acids. These ranges include the lengths tested in this study, but can be longer or shorter depending on the context. The target protein sequence is much longer than the sliding window. The sliding window is completely aligned (position for position) to the target protein sequence starting from the first amino acid on the target protein. At each position the absolute rounded difference in a biochemical metric is taken between the pair of amino acids at the particular point. Grantham polarity and Miyazawa hydrophobicity scales were used as the two metrics in the study. The information was sourced from expasyProtScale (https://web.expasy.org/protscale/).

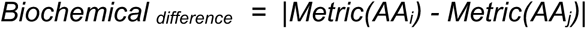

*Where*

*Metric: The value for a particular amino acid (AA) for a specific biochemical metric*

*i = The position on the sliding window*

*j = The position on the target protein,*

The rounded-off biochemical_difference_ is the encoding for the pair of amino acids at a given position on the two sequences. The encodings are concatenated sequentially to generate the interaction language. We add paddings in the end to ensure that the entire length of the target sequence is captured. Finally, the interaction language sequence is divided into k-sized subsequences, also called k-mers. The hyperparameter ‘k‘ is either tuned or chosen based on biological context. The k-mers are the input to the embedding architecture for generating the low-dimensional representations for each interaction. The doc2vec model is used for generating the interaction level embeddings. We used the doc2vec implementation from gensim^72^ (https://radimrehurek.com/gensim/) version 4.2.0. Two different architectures of the doc2vec model were used for the different biological interaction contexts. In the distributed memory (DM ) mode, a target biochemical k-mer is predicted using the embeddings of surrounding k-mers (also called the context window) along with the interaction embedding for a particular interaction. In the distributed bag of words (DBOW), the k-mers in a fixed length context window are predicted from the interaction embedding alone. The size of the context window, w can be adjusted in the embedding model. In both architectures, the k-mer embeddings and interaction embeddings are updated iteratively using stochastic gradient descent and back-propagation. In gensim, the DM hyperparameter is set as 1 for distributed memory architecture and 0 for distributed bag of words architecture. Besides the context length, other important doc2vec hyperparameters are the size of the vectors in the embedding matrix (dim), the minimum frequency of k-mers for inclusion in the model training (min_count) and the initial learning rate (alpha). After training the doc2vec model, the document embeddings are used as input features for training a classifier for interaction specific prediction tasks. We used xgboost’s XGBClassifier() (https://xgboost.readthedocs.io/en/stable/) function (version 1.6.1) and the sklearn’s LogisticRegression() (https://scikit-learn.org/stable/index.html) implementation as the classifiers. The important hyperparameters in xgboost are the number of boosting rounds (n_estimators), the maximum tree depth for the base learners (max_depth), and the learning rate for the boosting (learning_rate). The hyperparameters were tuned for the language, doc2vec and the classifier using the Bayesian optimization setting in weights and biases (WandB) (https://wandb.ai/site). The selection of the hyperparameters across the different models was selected in a supervised manner based on the performance of the classifier at a particular task. The evaluation setting for the hyperparameter optimization was 10 fold cross validation each for 10 replicates of the dataset (generated through changing the order of data points in the dataset). Either the mean of the area under the receiver operating characteristic curve (AUROC) or the F1 score was used for selecting the best model hyperparameters.

## Prediction of peptide-MHC interactions using SWING

Mass Spectrometry (MS) elution datasets were acquired from the publicly available HLA-specific NetMHCPan4.1 evaluation sets^73^, and NetMHCIIPan4.2 training sets^74^ for Class I and Class II respectively. The MHC sequences were acquired from the MHC Restriction Ontology (https://github.com/IEDB/MRO). The HLA molecules with available MS elution datasets were functionally clustered using MHCCluster 2.0^75^. For the Class I model datasets a subset of HLAs (*HLA*A02:02,* HLA*A02:11, HLA*A11:01, HLA*A25:01, *HLA*A32:01,* HLA*B07:02, HLA*B15:03, *HLA*B38:01, HLA*B40:01,*HLA*B40:02, HLA*C07:02, *HLA*C03:03, HLA*C05:01,* HLA*C08:02) were chosen. The subset was carefully curated to represent and test the diverse functional clusters identified through MHCCluster 2.0. Peptides larger than 12 amino acids were removed for all Class I sets. Similarly for the Class II model, a subset of single allele (SA) datasets (DRB1*01:01, DRB1*01:02, DRB1*03:01, DRB1*04:01, DRB1*04:04, DRB1*07:01, and DRB1*15:01) were selected.

We trained an XGboost model with the peptide-MHC embeddings as the features. For the Class I model, 8 HLA’s were chosen (HLA*A02:11, HLA*A11:01, HLA*A25:01, HLA*B07:02, HLA*B15:03, HLA*B40:02, HLA*C07:02, HLA*C08:02). The HLA’s were chosen from functionally distinct clusters such that each HLA group is similarly represented. 6 Class I (HLA*A02:02, HLA*A32:01, HLA*B40:02, HLA*C03:03, HLA*C05:01) datasets were chosen for validation. For the Class II model, the datasets were processed to remove any data that had more than one MHC molecule assigned to the entry. From the MHC molecules that were left, 5 (DRB1*01:01, DRB1*03:01, DRB1*04:01, DRB1*07:01, DRB1*15:01) were picked from different functional clusters for the training set, and *2* (DRB1:0102, DRB1:0404) were used as the validation. The training subset was chosen such that each DRB1 dataset was equally represented (∼30,000 peptides each). The language was generated using the peptide as the sliding window and the first 210 positions of MHC protein sequence as the target sequence. This region includes the Class I biologically relevant region (206 AA) of the alpha chains with 3 AA as padding. For Class II, the same cutoff on the beta chain was used to keep the models consistent. We used the Grantham polarity score to capture the difference between the amino acid pairs at each position of the two sequences. The training k-mers were shuffled for the cross prediction to prevent the model from overfitting. The doc2vec model (min_count =1, dm=0) parameters dim, w, alpha, and epochs along with the XGboost classifier parameters n_estimators, max_depth, and learning rate were tuned for optimal performance in predicting the interaction specific task. The final Class I model parameters are min_count=1, dm=0, k=7, dim=583, w=11, alpha=0.02349139979145104, epochs=13, n_estimators=232, max_depth=6, learning_rate=0.9402316101150048. The final Class II model parameters are min_count=1, dm=0, k=7, dim=146, w=12, alpha=0.03887032752085429, epochs=13, n_estimators=341, max_depth=9, learning_rate=0.6534638199102993.

First, the model was evaluated in a standard cross validation (SCV) setting. Here, ten fold cross validation was performed for each of the ten replicates of the dataset (generated by the shuffling of the order of the data points in the dataset) and the mean AUROC was reported for each time cross validation was performed. SCV was also performed for a permuted set where the labels were randomly generated with a 1:10 distribution of binders and non-binders, and the data shuffled in the same way as previously reported. SCV was also performed for a permuted set where the labels were randomly generated with a 1:10 distribution of binders and non-binders, and the data shuffled in the same way as previously reported. For visualization, the false positive rates and true positive rates at each threshold were obtained using the roc_curve() function in sklearn. Additionally, we performed interpolation using the numpy interp() function to calculate the mean of the AUROC and the standard deviation of the performance. The mean AUROC was plotted as a dashed line with shading to represent the 2 standard deviations as the performance bound. For testing the cross prediction performance of the Class I model, first, we chose the HLA alleles closer to the training dataset after the functional clustering. We performed 10 bootstraps on the validation dataset, keeping the label distribution intact for each subset of the validation set to mimic the entire dataset. The AUROC was calculated and visualized in the same way as described above. Second, we chose the validation set to comprise of HLA-alleles that were further away from the HLA alleles in the training set after the functional clustering. The steps were repeated for this validation set as performed for the preceding validation setting. The exact steps were repeated for testing the performance of the SWING Class II model. Due to the minimal number of SA Class II datasets available, 2 Class II datasets were chosen for the validation.

### Analyses for testing SWING’s performance on customized sequence inputs

We tested the effect of length on the MHC sequence on SWING’s performance. The longer MHC sequences encompass the full chain sequence and are between the range of 265-369. The biochemical metric was the Grantham polarity scale from expasyProtScale. The same evaluation setting was used for evaluating the Class I and II model with the longer sequence of the MHC molecules as in the previous section.

The second test was on the effect different biochemical scales have on SWING’s predictive power. We changed the scale from the Grantham polarity scale to the Miyazawa hydrophobicity scale, and the model was evaluated in the same way as described earlier.

Finally, the effect of the peptide length on SWING’s performance was checked. The effect of peptide length was tested on SWING’s performance by truncating the training and test set peptides to different sizes. The peptide sizes in the training set ranged the length of 13-21. Peptides larger than the cut-off size were truncated by taking the middle of the peptide. 16 different cut-off sizes (20,19,18,17,16,15,14,13,12,11,10,9,8,7,6,5) were used and the cross-prediction was performed as described and the AUC score was reported for 2 datasets (DRB1_0102, DRB1_0404). Additionally, the AUROC was reported for 20, 16, 12, and 8 cut-off size runs.

### Analyses to test SWING’s predictive performance between biologically distinct interactions

SWINGs performance for predicting between biologically different interaction spaces was evaluated. First, the performance of the Class I model was tested on 2 human Class II MHC (DRB1*01:02, DRB1*04:04) datasets using the cross-prediction method previously described without the bootstrapping. We compared the SWING Class I and Class II model performance as defined by AUC score with NetMHCPan4.1 (https://services.healthtech.dtu.dk/services/NetMHCpan-4.1/), NetMHCIIPan4.2 (https://services.healthtech.dtu.dk/services/NetMHCIIpan-4.2/), MixMHCPred 2.2 (https://github.com/GfellerLab/MixMHCpred), and MixMHC2Pred 2.0 (http://mixmhc2pred.gfellerlab.org/). Since the NetMHCPan4.1 method is not traditionally compatible with Class II data, the allele names (DRB1_0102, DRB1_0404) were manually added to the allele file and all relevant pseudo sequence files were copied from the NetMHCIIPan4.2 model. An empty space was provided for MixMHCPred 2.2 since it was not applied to the Class II data due to incompatibility with Class II alleles as well as peptides larger than 14. For the NetMHCpan-4.1,NetMHCIIpan-4.2, and MixMHC2Pred 2.0 models the “EL_score”, “Score”, and the negative of the “%Rank_best” columns were used as the predictions respectively.

Next, a Mixed Class model which consisted of 8 Class I (HLA*A02:11, HLA*A11:01, HLA*A25:01, HLA*B07:02, HLA*B15:03, HLA*B40:02, HLA*C07:02, HLA*C08:02) and ∼10,000 peptides from 2 Class II datasets (DRB1*04:01, DRB1*13:01) was trained. Hyperparameter optimization was performed and the final Mixed Class model parameters are min_count=1, dm=0, k=7, dim=74, w=12, alpha=0.03887032752085429, epochs=10, n_estimators=269, max_depth=9, learning_rate=0.6082359422582875. Model performance was evaluated through the SCV and cross-prediction on 3 Class I alleles (HLA-A02:02, HLA-B40:02, HLA-C05:01) and 2 Class II alleles (DRB1_0102, DRB1_0404). Lastly, a mouse Class II allele test set (H-2-IAb) was constructed from the NetMHCIIPan 4.2 training data and peptides available on IEDB, to test the performance of the Class II and Mixed Class models. The MHC sequence was retrieved from the Uniprot database. The AUROC was reported for each model.

### Profiling the I-Ek immunopeptidome

The protocol to profile the I-Ek immunopeptidome was adopted from Wan et al.^53^ with some modifications^76^. Kidneys from 20-week-old MRL/lpr mice were harvested and minced on a gentleMACS (Miltenyi Biotec) with Collagenase D (Roche) and DNase I (Sigma) in HBSS (Gibco)^77^. Cells were passed through a 70um filter to obtain a single cell suspension and RBC lysis was performed using RBC lysis buffer (Biolegend). The cells were counted and resuspended at 100x106 cells/mL in PBS (Corning). Cells were pelleted and incubated with lysis buffer (40mM MEGA 8 (Sigma-Aldrich), 40mM MEGA 9 (Sigma-Aldrich), 1mM PMSF (ThermoFisher Scientific), 0.2mM Iodoacetamide (Sigma-Aldrich), 20ug/ml Leupeptin (Millipore Sigma) and Roche Complete Mini Protease Cocktail (Roche)) for 1 hour at 4°C on a rocking platform. The lysate was cleared by centrifugation at 20,000g for 30 min at 4°C. Anti-MHC Class II I-Ek antibody [14-4-4S] (Abcam) was covalently linked to sepharose G resin (Cytiva) by incubation for 30 min in cold room on rocker. The cleared lysate was transferred to a tube containing sepharose G coupled to antibody and incubated at 4°C overnight on a rocker. The I-Ek sepharose lysate was poured in a Poly-Prep Chromatography Column (Biorad) and gradient washes were performed with the following buffers serially: 10ml 150mM NaCl, 20mM Tris pH7.4, 10ml 400mM NaCl, 20mM Tris pH7.4, 10ml 150mM NaCl, 20mM Tris pH7.4, 10ml 20mM Tris pH8.0. The peptides were eluted in 10% acetic acid and sent for mass spectrometry to MS Bioworks.Peptides (50% per sample) were acidified, then concentrated and desalted using solid-phase extraction (SPE) with Waters µHLB C18 plate. Peptides were loaded directly and eluted using 30/70 acetonitrile/water (0.1% TFA). Eluted peptides were lyophilized and reconstituted in 0.1% TFA. Peptides (100%) were analyzed by nano LC/MS/MS using a Waters NanoAcquity system interfaced to a ThermoFisher Fusion Lumos mass spectrometer. Peptides were loaded on a trapping column and eluted over a 75µm analytical column at 350nL/min; both columns were packed with Luna C18 resin (Phenomenex). A 2h gradient was employed. The mass spectrometer was operated using a custom data dependent method, with MS performed in the Orbitrap at 60,000 FWHM resolution and sequential MS/MS performed using high resolution CID and EThcD in the Orbitrap at 15,000 FWHM resolution. All MS data were acquired from m/z 300-1600. A 3s cycle time was employed for all steps.Peptides were analyzed using PEAKS software with the following parameters, Enzyme: None, Database: Swissprot Mouse, Fixed modification: None, Variable modifications: Oxidation (M), Acetyl (Protein N-terminus), Carbamidomethyl (C), Mass values: Monoisotopic, Peptide Mass Tolerance: 10 ppm, Fragment Mass Tolerance: 0.02 Da, Max Missed Cleavages: N/A, PSM FDR: 1%, Chimeric peptide: TRUE.

### Evaluating the performance of SWING on the de novo datasets

The accuracy of multiple pMHC prediction tools were evaluated using the experimentally generated H-2-IEk dataset. The MHC sequence for the allele was retrieved from Uniprot. H-2-IEk binders between the length of 9-25 with standard amino acids and no modifications were considered. This subset was evaluated using the 3 SWING models as well as 2 benchmarking methods, NetMHCIIPan4.2 and MixMHC2Pred 2.0. For the SWING models, we predicted the probability that the peptides are interactors of H-2-IEk. A cut-off threshold was identified using the highest geometric mean of the false positive rate and true positive rate from the SCV of the Class I (0.0012), Class II (0.008), and Mixed Class (0.002) models. Predicted probabilities that were larger than the cut off threshold were considered binders. For NetMHCIIPan4.2, the web server was used. Peptides were split into different groups by size and each group was run through with the H-2-IEk allele as the chosen sequence and default parameters. For the MixMHC2Pred 2.0, only peptides of length 12-21 were provided due to the limitations imposed on peptide length by the method. Peptides with a rank <30 were considered interactors. Accuracy was used as the performance metric for all models. An accuracy was calculated for each peptide length for the range of 9-25 and plotted. Additionally, a total accuracy score was calculated. Since MixMHC2Pred 2.0 is only able to calculate a score for peptides 12-21, this range was used for the total accuracy calculation. We repeated this procedure with a second, previously published dataset for the H-2-IAg7 allele (Zdinak Nature Methods In Press). Since NetMHCIIPan 2.0 does not have the H-2-IAg7 allele in the server, the sequence was provided for the beta chain.

### Analyses to test SWING’s performance on predicting interaction specific effects of missense mutations

The dataset for predicting the impact of missense mutations on protein interactions was sourced from two papers studying the impact of population variants on interactions across the allele frequency spectrum^61^ and the effect of disease associated variants on the interactions respectively^63^. The ENTREZ gene IDs were mapped to the respective uniprot IDs and the dataset was filtered for hetero-dimers. The uniprot database was used for mapping the proteins to the respective amino acid sequences. The resulting dataset consisted of 3,386 missense mutations tested across 3,422 PPIs. With respect to the training procedure, the reported variable was considered to be the disruption of the PPI by the missense mutation tested through a yeast-2- hybrid assay for both the datasets.

For the task of predicting the impact of missense mutations on PPIs, the window is selected from the protein sequence with the mutation. Particularly, L positions on both sides of the mutation affected site are used to select the sliding window where L is a tunable parameter. The language generation task is the same as described above. The string generated from this process is split into k-mers, where k is another tunable parameter. We used a window size (L) of 1 and k-mers of size 7.

The embeddings generated for the protein-perturbed peptide pair were fed as the features to the classifier to predict the disruption of an interaction. The training datasets had population variants across the allele frequency spectrum or are associated with Mendelian diseases. The alpha (learning rate) of the doc2vec model (fixed dm=1, dimension=128, epochs=200, and min_count=1) and n_estimators, max_depth, and learning_rate of the XGboost classifier are tuned for optimal performance of the classifier using a Bayesian approach through the framework provided in Weights and Biases - WandB (wandb.ai). Doc2Vec parameters included an alpha of 0.0223, 200 epochs, vector size of 128, and used distributed memory (dm=1). XGBoost had 225 estimators, max depth of 6, and learning rate of 0.186.

The standard cross validation was performed on three different models, a SWING model trained on just mendelian disease associated variants, model trained with population variants and a model trained on both the datasets. The evaluation framework was the same as described in the sections above.

We constructed a “leave cluster out” framework for a more rigorous evaluation of the model. Each amino acid in the sequence of the interaction partner protein of the mutated protein was originally encoded with a number between 0 to 20. Subsequently, we performed KMeans clustering with k=10 on the encoded sequences to assign clusters using sklearn. We then applied sklean’s PCA function to the encodings to visualize them in a 2D space, coloring based on cluster assignment. Sets of two clusters were considered a fold and the two closest positionally adjacent clusters were dropped from the training set for complete exclusivity between the training and testing sets. The AUROC for each test cluster was reported individually.

AlphaMissense^59^ , ESM1b^60^ and EVE^58^ mutation effect prediction models (LLM tools) were chosen for benchmarking. We checked whether there is a difference in the pathogenicity scores for the interface and the non-interface localized variants. We selected only those mutations whose interaction perturbation profiles were present in the training dataset curated for SWING and the corresponding co-crystal structure for the interaction were present in the PDB. Additionally, These were all ClinVar variants and were reported in Cheng et al^59^. The final evaluation set was the intersection of the ClinVar variants at the interface of structurally resolved interactions and missense mutations which disrupted the particular interactions. A Mann Whitney-U test was done to test the difference in the scores of interface versus non-interface localized variants. The p-values were transformed to negative of log_10_ and reported. The LLM tools scores were tested for predictive power on interaction specific effects by ascribing the scores with each interaction that has been tested for the particular variant. The AUROC score was reported for each method and compared to the standard cross validation AUROC from SWING.

The SWING model that had been trained on mendelian and population variant interaction effects datasets was used to predict the probability of disruption by ClinVar variants present in the AlphaMissense study^59^ and absent from SWINGs training dataset. The threshold for SWING classifying a variant as highly interaction disruptive was 0.9 and the threshold for alphamissense to classify a variant as benign was 0.5. Clinvar pathogenicity annotations (https://www.ncbi.nlm.nih.gov/clinvar/) were used to determine the literature reported pathogenic status of the variant.

### Testing the effect of the hydrophobicity scale on the prediction performance for predicting the effect of mutation on PPIs

The interaction language was encoded using the Miyazawa hydrophobicity from expasyProtScale. We used the same doc2vec implementation from gensim^72^ (https://radimrehurek.com/gensim/) version 4.2.0 and XGBClassifier() from xgboost (https://xgboost.readthedocs.io/en/stable/) version 1.6.1 implementation as the classifier. The hyperparameters tuned from WandB’s Bayesian optimization setting (https://wandb.ai/site) were the same as mentioned in a preceding section (Language generation: k=7, L=1; doc2vec: alpha=0.0223, epochs=200, vector_size=128, dm=1; XGBoost: n_estimators=225, max_depth=6, learning_rate=0.186).

### Predicting missense mutation mediated interaction disruption using logistic regression

We also wanted to show that a simpler model was able to capture signal in predicting the impact of missense mutations on PPIs using the interaction language. We used the same doc2vec implementation from gensim^72^ (https://radimrehurek.com/gensim/) version 4.2.0 but replaced the XGBoost Classifier with sklearn’s LogisticRegression() (https://scikit-learn.org/stable/index.html) version 1.1.1 implementation as the classifier. The hyperparameters used for language generation and doc2vec were the same as Figure 5 (k=7, L=1, alpha=0.0223, epochs=200, vector_size=128, dm=1), but we used WandB’s Bayesian optimization setting (https://wandb.ai/site) to tune LogisticRegression() on C and l1_ratio, fixing solver to saga and penalty to elasticnet. The tuned LogisticRegression() parameters were l1_ratio=0.06056 and C=0.6012. While this simpler model found a signal, we found XGBoost() performed better at the missense mutation disruption prediction task.

### Predicting missense mutation mediated interaction disruption using a multilayer perceptron

Using the same doc2vec implementation from gensim^72^ version 4.2.0, we implemented sklearn’s MLPClassifier() (https://scikit-learn.org/stable/index.html) version 1.1.1 as our classifier. The hyperparameters used for language generation and doc2vec were again the same as Figure 5 (k=7, L=1, alpha=0.0223, epochs=200, vector_size=128, dm=1), but we used WandB’s Bayesian optimization setting (https://wandb.ai/site) to tune MLPClassifier() on activation, alpha, hidden_layer_sizes, and learning_rate with early_stopping=True. The tuned MLPClassifier() parameters were activation=’tanh’, alpha=1.76, hidden_layer_sizes=(226,), and learning_rate=’constant’.

## Data and Code Availability

All datasets, code and documentation are available at https://github.com/jishnu-lab/SWING.

## Author Contributions

J.D., A.J. and P.C. conceived of the project with conceptual inputs from D.K.. J.D. and A.J. supervised all aspects of the study. J.D. designed all computational analyses which were implemented by A.O., J.S., P.C. A.R. and J.R. also helped with the implementation. A.O., J.S. and P.C. contributed equally and have the right to list their names first in their curriculum vitae. I.N. and K.N. performed the benchmarking of SWING against existing methods. M.S. and J.T. provided the necessary reagents and mouse models for SLE and T1D immunopeptidome discovery. A.J. designed the immunopeptidome experiments and S.A performed the respective experiments. A.O., J.S., P.C. and J.D. wrote the manuscript with input from all authors.

## Conflict of Interest

The authors declare no conflict of interest.

## Acknowledgements

J.D. was supported in part by NIAID DP2AI164325, NIAID R01AI170108 and NHGRI U01HG012041. This research was supported in part by the University of Pittsburgh Center for Research Computing through the resources provided.

## Supplementary Figure Legends

**Supplementary Figure 1:**

Peptide length distribution of the non-interacting peptides in the Class II datasets defined by percentage. Magenta, training set; blue, DRB1_0102 validation set; orange, DRB1_0404 validation set.

**Supplementary Figure 2:**

Prediction performance in a standard cross validation setting of SWING trained for predicting the interaction perturbation effect by mendelian missense mutations (pink star), population variants (purple triangle), and a mixed set of mendelian missense mutations and population variants using hydrophobic scale. Blue, Interaction perturbation effects; Red, Permuted classifier; Green, Perfect classifier; Grey, Random Classifier.

**Supplementary Figure 3:**

Prediction performance in a standard cross validation setting of SWING trained for predicting the interaction perturbation effect by mendelian missense mutations (pink star), population variants (purple triangle), and a mixed set of mendelian missense mutations and population variants using a logistic regression classifier. Blue, Interaction perturbation effects; Red, Permuted classifier; Green, Perfect classifier; Grey, Random Classifier.

**Supplementary Figure 4:**

Prediction performance in a standard cross validation setting of SWING trained for predicting the interaction perturbation effect by mendelian missense mutations (pink star), population variants (purple triangle), and a mixed set of mendelian missense mutations and population variants using a multilayer perceptron classifier. Blue, Interaction perturbation effects; Red, Permuted classifier; Green, Perfect classifier; Grey, Random Classifier.

**Supplementary Table 1:**

Scores for SWING Class I, SWING Class II, netMHCpan 4.1, mixMHCpred 2.0, netMHCIIpan 4.2, and mixMHC2pred 2.0 models for predicting Class II pMHC interactions for DRB1_0102 and DRB1_0404 validation sets.

**Supplementary Table 2:**

Label scores for H-2-IEk and H-2-IAg7 interacting peptides for different SWING models, netMHCIIPan 4.2, and MixMHC2Pred 2.0.

**Supplementary Table 3:**

Interface variants from ClinVar with interaction disruption scores and AlphaMissense pathogenicity scores

## Notes

### Competing Interest Statement

The authors have declared no competing interest.

