## Supplementary figures and images for "Sliding Window INteraction Grammar (SWING): a generalized interaction language model for peptide and protein interactions"

### Supplementary Figures

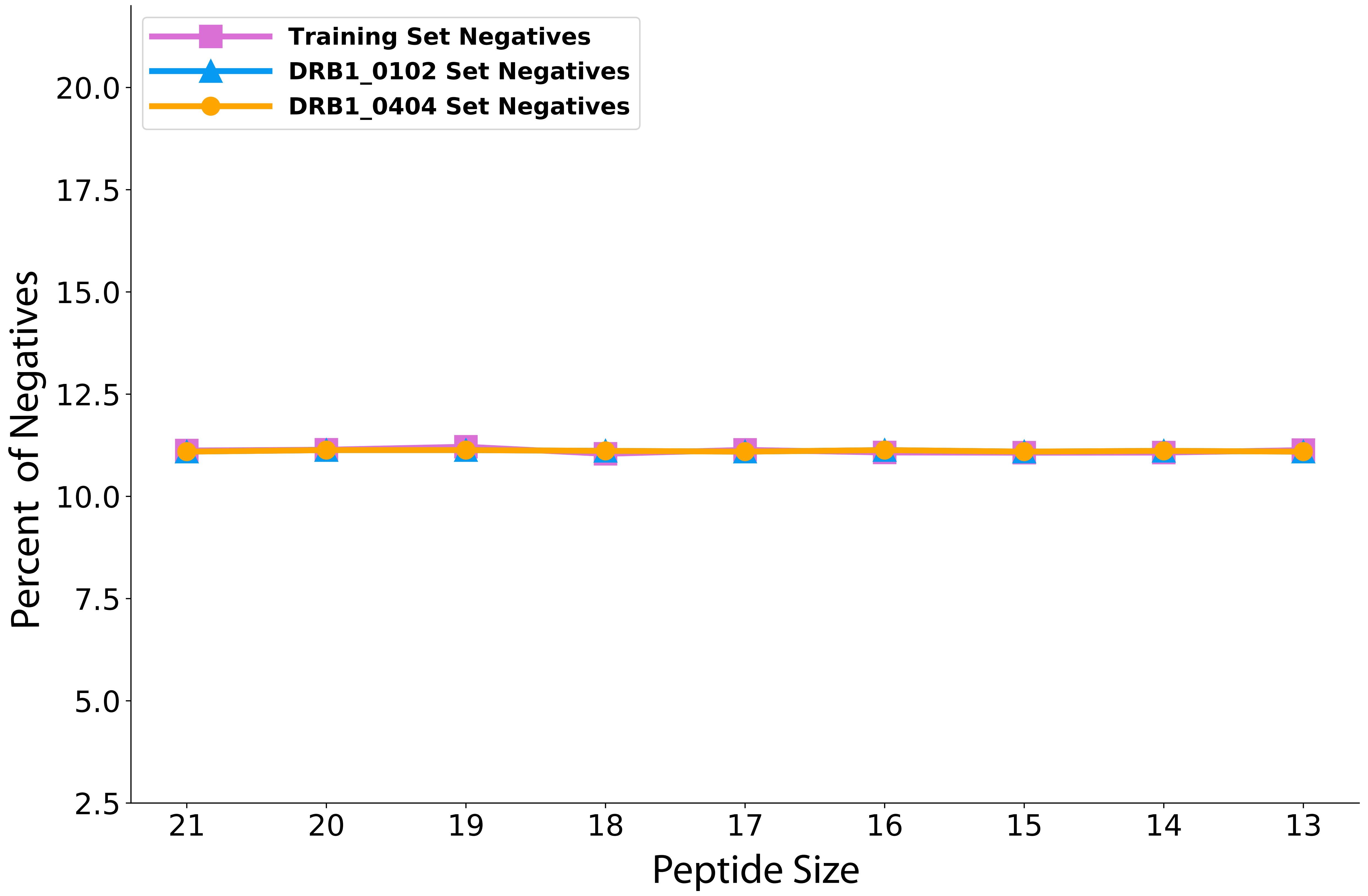

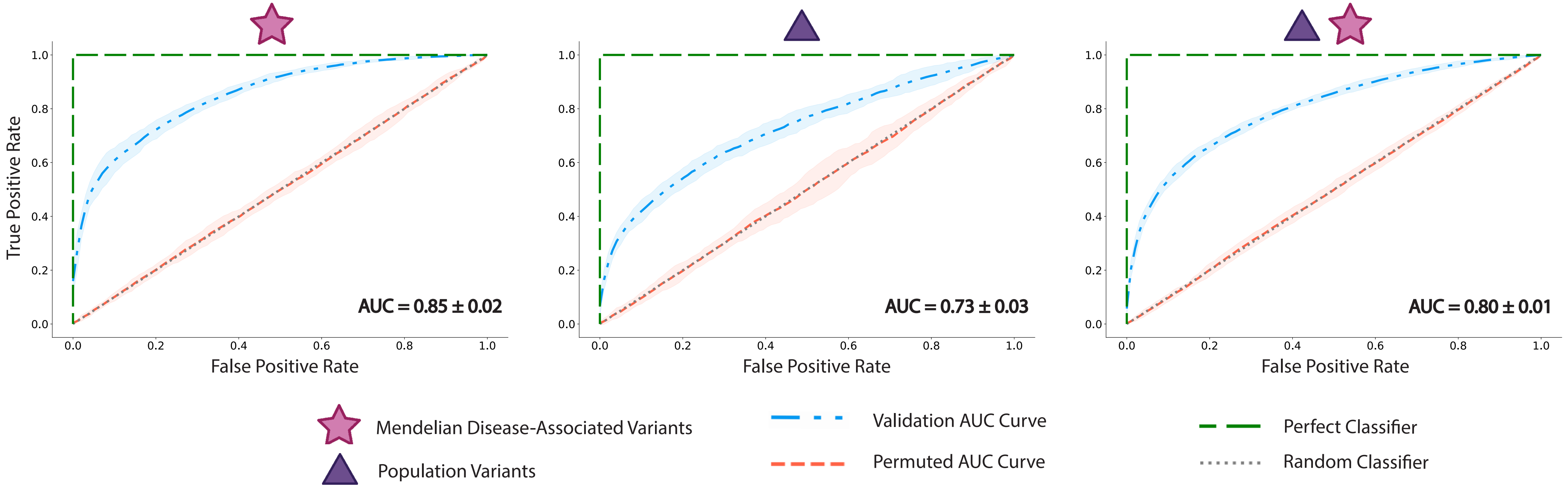

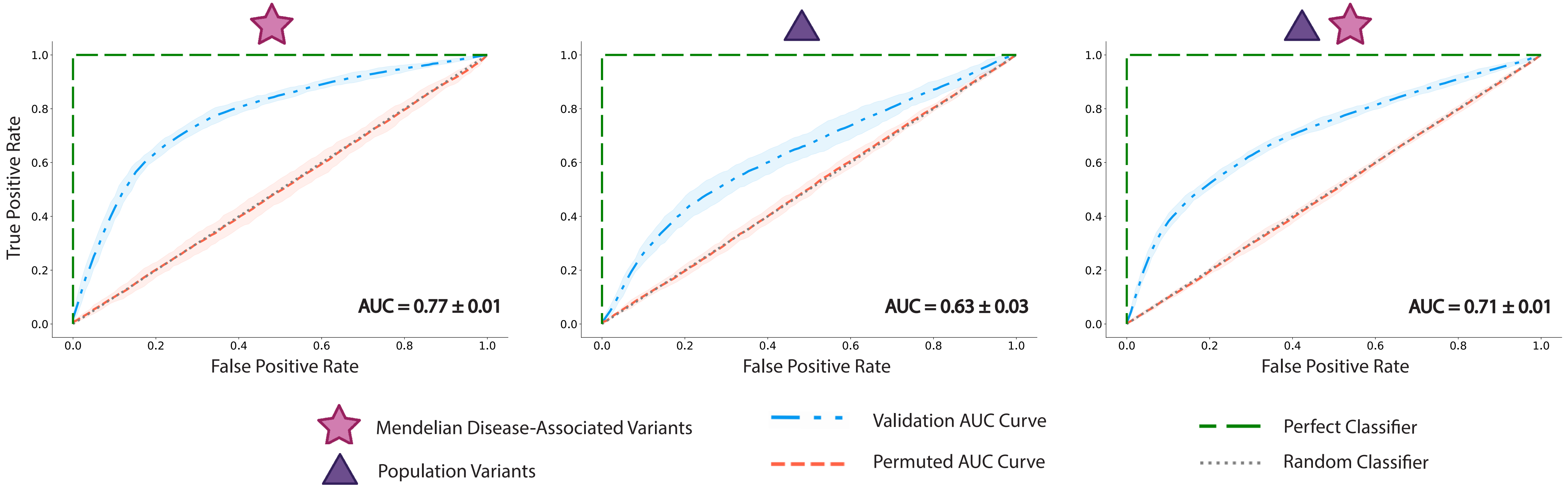

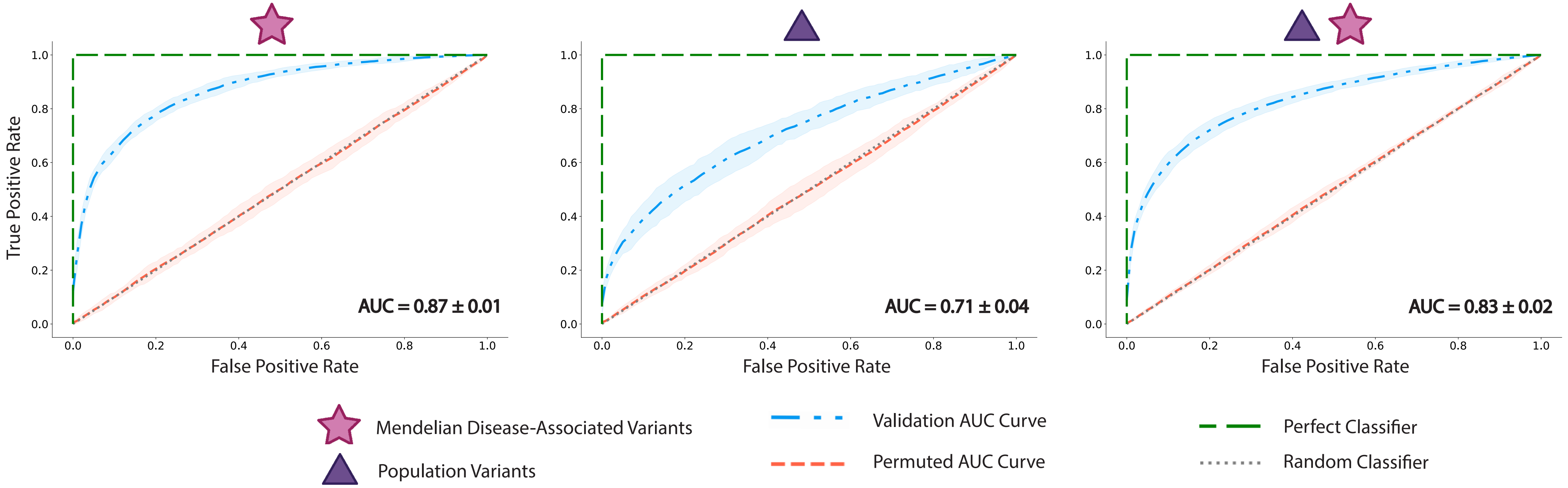
